# Circuits for integrating learnt and innate valences in the fly brain

**DOI:** 10.1101/2020.04.23.058339

**Authors:** Claire Eschbach, Akira Fushiki, Michael Winding, Bruno Afonso, Ingrid V Andrade, Benjamin T Cocanougher, Katharina Eichler, Ruben Gepner, Guangwei Si, Javier Valdes-Aleman, Marc Gershow, Gregory SXE Jefferis, James W Truman, Richard D Fetter, Aravinthan Samuel, Albert Cardona, Marta Zlatic

## Abstract

Animal behavior is shaped both by evolution and by individual experience. In many species parallel brain pathways are thought to encode innate and learnt behavior drives and as a result may link the same sensory cue to different actions if innate and learnt drives are in opposition. How these opposing drives are integrated into a single coherent action is not well understood. In insects, the Mushroom Body Output Neurons (MBONs) and the Lateral Horn Neurons (LHNs) are thought to provide the learnt and innate drives, respectively. However their patterns of convergence and the mechanisms by which their outputs are used to select actions are not well understood. We used electron microscopy reconstruction to comprehensively map the downstream targets of all MBONs in *Drosophila* larva and characterise their patterns of convergence with LHNs. We discovered convergence neurons that receive direct input from MBONs and LHNs and compare opposite behaviour drives. Functional imaging and optogenetic manipulation suggest these convergence neurons compute the overall predicted value of approaching or avoiding an odor and mediate action selection. Our study describes the circuit mechanisms allowing integration of opposing drives from parallel olfactory pathways.

## Introduction

Selecting appropriate actions in response to sensory stimuli is a major brain function. To achieve this, brains must transform complex representations of sensory stimuli into representations of valences (attractiveness or aversiveness) that can be used to drive actions (Pearson et al., 2014). Many sensory stimuli have innate valences, acquired through evolution: some stimuli are innately attractive and others are innately repulsive (Q. Li & Liberles, 2015; Reisenman et al., 2016). However, to behave adaptively in an ever-changing environment, animals are also able to learn new valences for stimuli (Q. Li & Liberles, 2015). These learnt valences can be in conflict with innate ones. For example, repeated association of an innately attractive odor with punishment (*e.g.* pain, or illness) allows a switch from innate attraction to learnt aversion of the same odor (Garcia et al., 1983; Pauls et al., 2010; Tully & Quinn, 1985). The innate and learnt valences are thought to be encoded in distinct brain areas in both vertebrates (Choi et al., 2011; Q. Li & Liberles, 2015; Sosulski et al., 2011) and invertebrates (Q. Li & Liberles, 2015; Marin et al., 2002). In mammals, the olfactory projection neurons (mitral cells) send divergent projections to two parallel higher-order centers, the olfactory amygdala and the piriform cortex, implicated in innate and learnt behaviors, respectively (Choi et al., 2011; Q. Li & Liberles, 2015; Root et al., 2014; Sosulski et al., 2011). Likewise, in insects the olfactory projection neurons send divergent projections to the lateral horn (LH) and the mushroom body (MB, (Eichler et al., 2017; Gerber & Stocker, 2007; Jeanne et al., 2018; Marin et al., 2002; Wong et al., 2002), implicated in innate and learnt behaviors, respectively (Yoshinori Aso, Hattori, et al., 2014; Yoshinori Aso, Sitaraman, et al., 2014; Dolan et al., 2019; Heimbeck et al., 2001; Heisenberg, 2003; Q. Li & Liberles, 2015; Parnas et al., 2013; Turner et al., 2008). Thus, two distinct olfactory structures output valence signals that can be used for an odor response, but the way in which these signals are used to produce a coherent behavioral choice is still an open question. For example, how are conflicting valence signals resolved? Do opposing drives for behavior converge and get integrated, or do they remain in competition (Pearson et al., 2014)?

A major obstacle to addressing these questions has been the lack of comprehensive synaptic-resolution maps of the patterns of convergence between neurons that represent innate and learnt valences. Another obstacle has been the inability to causally relate specific circuit elements to their function. Here, we were able to overcome these obstacles by using the tractable genetic model system of *Drosophila melanogaster* larva. In this system, we could combine: i) large-scale electron microscopy reconstruction of neural circuits due to the relatively small size of its brain (Jovanic et al., 2016; Ohyama et al., 2015); ii) targeted manipulation of uniquely identified neuron types (Jovanic et al., 2016; Ohyama et al., 2015; Tastekin et al., 2018), iii) and functional imaging of neural activity.

Previous studies in *Drosophila* have characterized all the components of the MB network and their roles in memory formation and expression. The MB consists of a set of parallel fiber neurons, the Kenyon cells (KCs), that sparsely encode sensory inputs coming from olfactory and other projection neurons (PNs, (Gerber & Stocker, 2007; Hige, Aso, Rubin, et al., 2015; Honegger et al., 2011; Lin et al., 2014; Papadopoulou et al., 2011). KC axons are tiled into distinct compartments by terminals of modulatory neurons, mainly dopaminergic (DANs, (Yoshinori Aso, Hattori, et al., 2014; Eichler et al., 2017; Mao & Davis, 2009; Takemura et al., 2017)). DANs carry information about positive and negative reinforcement and provide teaching signals for memory formation (Yoshinori Aso, Hattori, et al., 2014; Y. Aso & Rubin, 2016; Eschbach et al., 2020; C. Liu et al., 2012). In each compartment, DANs synapse onto KCs and onto the dendrites of compartment-specific MB output neurons (MBONs; (Eichler et al., 2017; Takemura et al., 2017)). In the adult, individual MBONs have been shown to promote approach or avoidance (Yoshinori Aso, Sitaraman, et al., 2014; Bouzaiane et al., 2015; David Owald & Waddell, 2015). Pairing of an odor with a DAN has also been shown to selectively depress the conditioned odor drive to MBONs in that compartment (Hige, Aso, Modi, et al., 2015). Prior to learning, aversive and appetitive MBONs are thought to receive similar odor drive. Aversive and appetitive learning depress the odor drive to appetitive and aversive MBONs, respectively (Yoshinori Aso, Sitaraman, et al., 2014; David Owald & Waddell, 2015). Learnt valence of stimuli is therefore thought to be encoded as a skew in the activity of the population of MBONs. However, the way in which the learnt valences are read out by the networks downstream of MBONs and used to select actions is mostly unknown. Similarly, the way in which innate and learnt valences are integrated is poorly understood. In principle, MBONs could directly modify innate valences by directly synapsing onto LH neurons (LHNs), LHNs could synapse directly onto MBONs, or LHNs and MBONs could converge onto downstream neurons. Recent studies in *Drosophila* adult have uncovered one pathway of integration involving direct input from MBONs onto LHNs (Dolan et al., 2018, 2019; Lerner et al., 2020). However, whether this is the only mechanism, or whether additional patterns of convergence exist is unclear, because a comprehensive synaptic-resolution characterisation of the structural patterns of convergence between MBONs and LHNs was lacking.

Here, we first confirmed that larvae can switch innate odor attraction to learnt odor avoidance, after an innately attractive odor is paired with a noxious stimulus. We then determined which larval MBONs promote approach or avoidance when optogenetically activated. Next, we exhaustively reconstructed all neurons postsynaptic to all MBONs, as well as all LHNs in an innately attractive pathway. Together, these reconstructions provide a comprehensive view of the structural patterns of convergence between brain areas that encode innate and learnt valences. They revealed that some MBONs directly synapse onto LHNs as previously shown in the adult. However, we also identified two novel patterns of convergence: i) some LHNs directly synapse onto some MBONs, and ii) MBONs and LHNs converge onto downstream “convergence neurons (CNs)”. We found that some CNs compare excitatory and inhibitory input from MBONs and LHNs that promote opposite actions. We found that an increase in the activity of these neurons drives opposite actions. Together, our studies suggest these CNs integrate learnt and innate valences to compute an overall predicted value associated with approaching or avoiding an odor and promote actions consistent with the predictions they encode. These studies provide mechanistic insight into how conflict between opposing valences can be resolved by a population of integrative neurons.

## Results

### Larvae switch from innate odor approach to learnt odor avoidance by modulating turning

*Drosophila* larvae are innately attracted by most volatile molecules (Fishilevich & Vosshall, 2005; Kreher et al., 2008; Mathew et al., 2013), and repelled by a few, such as CO2 (Gershow et al., 2012). Larvae navigate gradients of innately attractive or repulsive odors via klinotaxis (Gepner et al., 2015; Gershow et al., 2012; Gomez-Marin & Louis, 2012; Schulze et al., 2015): larval motion alternates between turns and crawls. The probability and direction of turns versus the probability and speed of crawling events are modulated by changes in odor intensity. Larvae display odor attraction when the coupling between odor intensity change and turn probability is negative; when positive, they display odor avoidance (Gepner et al., 2015; Gershow et al., 2012; Gomez-Marin & Louis, 2012; Schulze et al., 2015). In other words, larvae approach innately attractive odors using as follows: i) When crawling towards the attractive odor source they sense an increase in the concentration of the odor so they repress turning (and promote crawling); ii) when crawling away from the attractive odor source they sense a decrease in the concentration of the odor so they promote turning (and repress crawling). They avoid innately aversive odors by doing the opposite. Associative learning has been shown to modify the coupling between odor intensity change and turn probability (Paisios et al., 2017; Michael Schleyer et al., 2015). Aversive learning (pairing an odor with punishment or absence of a reward) has been shown to switch the coupling (Paisios et al., 2017). Here we report similar findings using optogenetic punishment. We compared larvae that received an innately attractive odor, ethyl acetate (Gershow et al., 2012; Kreher et al., 2008), paired with the activation of the nociceptive Basin interneurons (Ohyama et al., 2015) to larvae that received unpaired presentation of the two stimuli (Fig. 1a-c). As expected, we found that pairing the odor with optogenetic punishment switched innate odor attraction to learnt avoidance. This involved a switch from negative to positive coupling between odor intensity change and turn probability: the paired group turned more when crawling up the odor gradient, than down the odor gradient, whereas the unpaired group larvae did the opposite (Fig. 1b-c). The rest of this article will focus on investigating the neural basis of the switch in coupling between odor intensity change and turning following aversive training.

**Figure 1:**
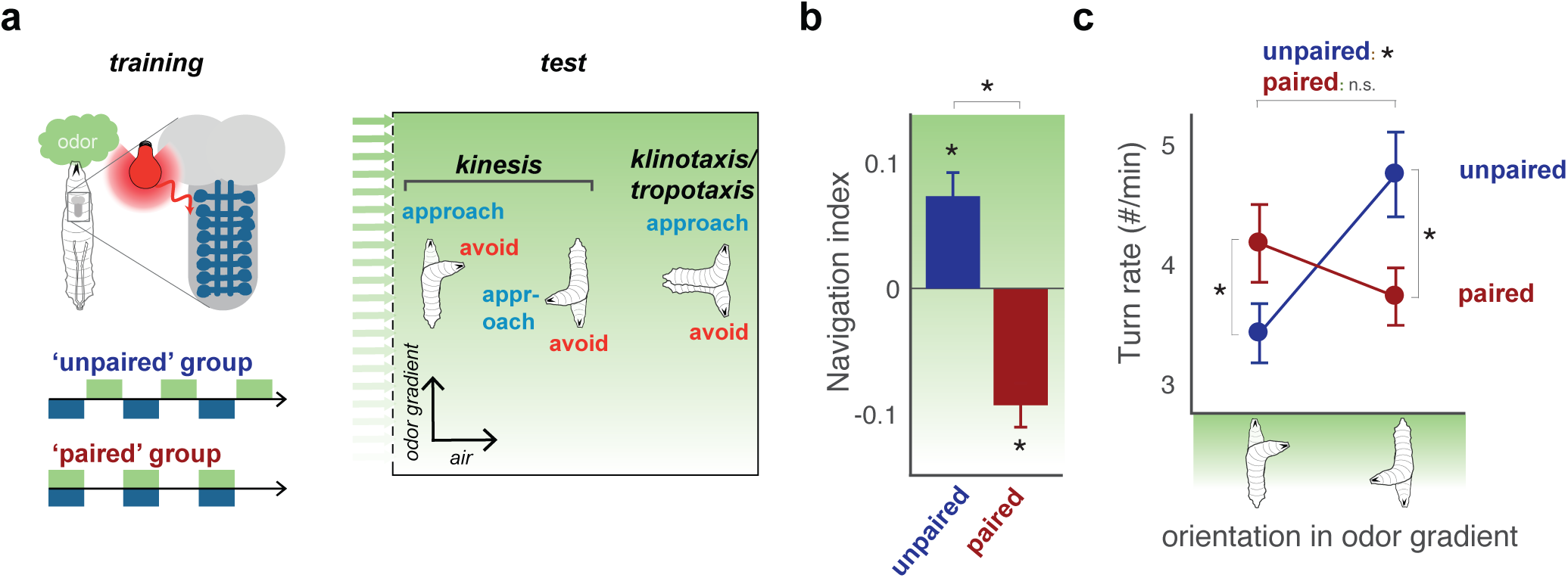
Aversive learning induces a switch in odor navigation strategy. **a.** The behavior of larvae in a linear gradient of odor (ethyl acetate) is recorded after the animals have undergone odor presentation intercalated (unpaired training protocol) or coincident (paired training protocol) with fictive punishment (optogenetic activation of nociceptive neurons). Larvae navigate via a strategy involving kinesis, where they modulate their turn rate over time and in response to different conditions, and klinotaxis/tropotaxis, where they choose turn side. Here, we record turn rate as a function of the orientation of the larva in the odor gradient (**c**). **b.** Navigational indices are obtained by dividing the mean velocity in the direction of the gradient by the mean crawling speed. The positive index after the unpaired protocol indicates that larvae approach the odor, the negative index after the paired protocol indicates that larvae avoid the odor. **c.** Turn rate versus orientation in the gradient. After unpaired and paired protocol, respectively, larvae exhibit positive and negative kinesis. Thus, after aversive learning with the odor, the larvae avoid the odor by altering the correspondence between turn rate and orientation in the odor gradient. Values are mean ± s.e.m. *: p < 0.05 in a Welch *Z*-test, N = 10 repeats.

### Identification of approach- and avoidance-promoting MBONs

Since olfactory learning has been found to modify the strength of KC-to-MBON synapses in the adult (Hige, Aso, Modi, et al., 2015), we first investigated how individual MBONs influence approach and avoidance, and in particular turning. We generated Split-GAL4 lines (Pfeiffer et al., 2010) that allowed us to drive expression of the red-shifted channelrhodopsin CsChrimson (Klapoetke et al., 2014) in a single or in a couple of indistinguishable MBONs per brain hemisphere (Extended Data Fig. 1). We then monitored behavioral responses to optogenetic activation of individual MBON types (Fig. 2a-g). Activating some MBONs had no effect on behavior (Extended Data Fig. 2). Most MBONs tested fell into one of two opposing categories in terms of their effect on behaviour.

**Figure 2:**
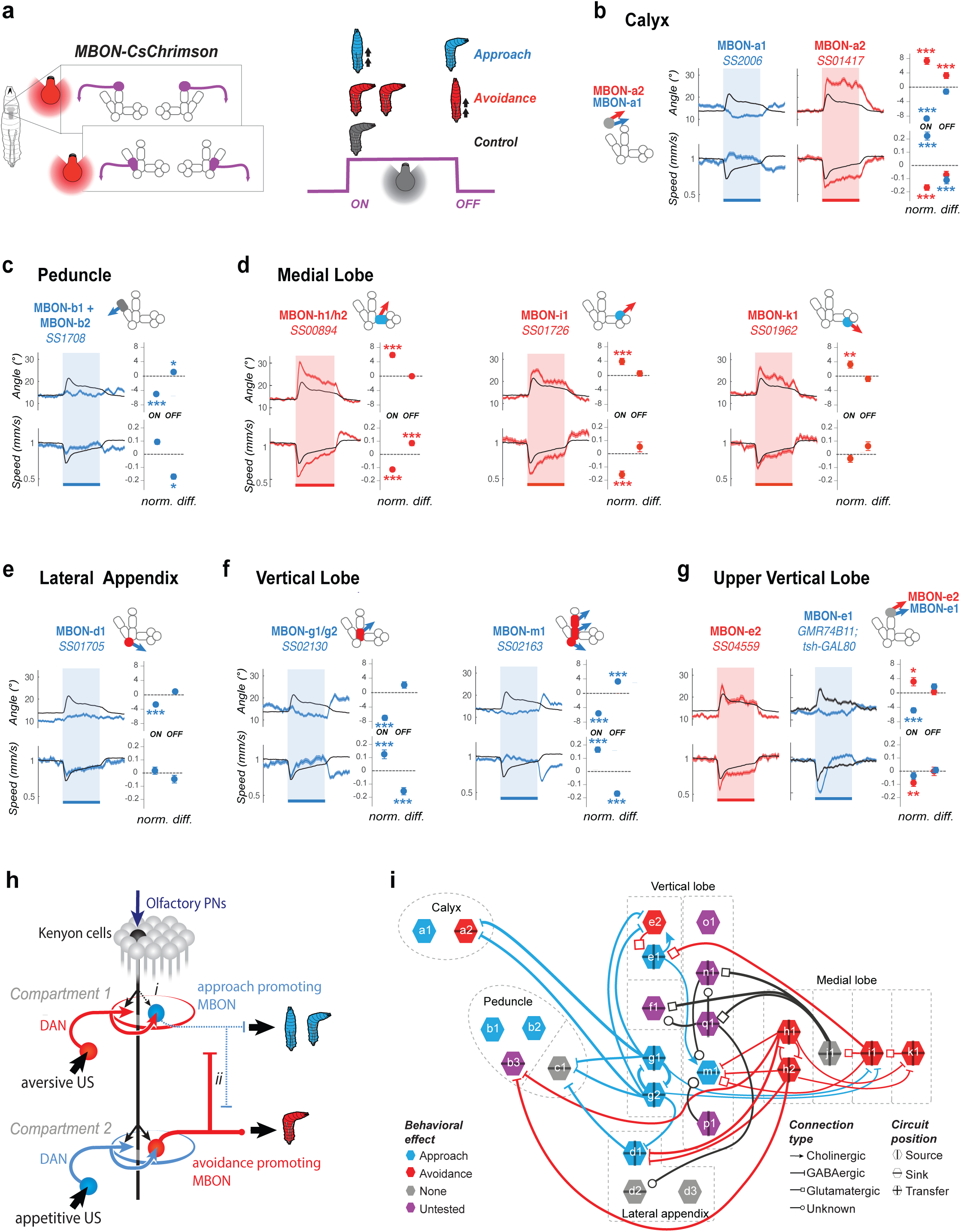
MBONs can promote odor approach or avoidance and are hierarchically organized. **a.** We recorded the behavior of larvae expressing CsChrimson in one or two pairs of MBONs and classified their responses to a 15-sec red-light stimulation as approach-like (*blue larva*), avoidance-like (*red larva*), or neutral by comparing them to the responses of empty driver line control animals. Approach-like responses are characterized by significantly decreased turning and/or increased crawling speed in response to an increase in activity (at the onset of optogenetic activation) compared to controls, and/or increased turning and decreased crawling in response to a decrease in activity (at the offset of optogenetic activation). Avoidance-like responses are characterized by the reverse responses. (**b.-g.**) Behavioral response to optogenetic activation of one to two pairs of MBONs. The GAL4 or Split-GAL4 lines used are indicated in italic and their expression patterns are visible in Extended Data Figure 1. The lines that drive expression in MBONs with no visible activation phenotypes are shown in Extended Data Figure 2. The schematics depict the compartments where the MBONs extend their dendrites, filled with color indicating which type of memory this compartment can form (Eschbach *et al*., 2020): appetitive short-term (*blue*), aversive short-term (*red*), or unknown (*grey*). Left plots show mean turn angle (*top*) and mean crawling speed (normalized to baseline, *bottom*) as a function of time. Colored bars and shades indicate the period of optogenetic activation. Values are mean +/- s.e.m. Right plots show the difference between the experimental (red or blue dots with error bars) and control (dotted line at 0) lines in the value of the same parameters averaged over a time window at the onset (0-5 sec. after light on, normalized to baseline before light) and offset of optogenetic stimulation (2-7 sec. after light off, normalized to baseline during light). Note that the control animals (black curve in **b-g**) displays a slightly aversive response to the onset of red light used for optogenetic activation. The control is the empty line *y w;attP40;attP2* crossed to *UAS-CsChrimson* (N_exp_=343,N_larvae_>10000) for all lines except for MBON-e1 (**g**) for which the control line is *yw;;attP2* crossed to *UAS-CsChrimson; tsh-GAL80*. Plots are mean +/- s.e.m. *: p<0.005, **: p<0.001, ***: p<0.0001 (Welch’s Z test). **b.** Activating the two calyx-MBONs induced opposite responses: approach and avoidance, for MBON-a1 (N_exp_=7, N_larvae_≈250) and MBON-a2 (N_exp_=7, N_larvae_≈280), respectively. **c.** Activating peduncle-MBONs, MBON-b1 and MBON-b2 together (N_exp_=8, N_larvae_≈340) induced approach. MBON-c1 did not have a significant effect on behavior (Extended Data Figure 2). **d.** Activating medial lobe MBONs induced avoidance: MBON-h1/h2 (N_exp_=10, N_larvae_≈450), MBON-i1 (N_exp_=6, N_larvae_≈240) and MBON-k1 (N_exp_=6, N_larvae_≈210). No effect was observed for MBON-j1 activation (Extended Data Figure 2). **e.** Activating the lateral appendix-MBON-d1 (N_exp_=9, N_larvae_≈250) induced approach, whereas no effect was observed for MBON-d2 and MBON-d3 activation (Extended Data Figure 2). **f.** Activating the vertical lobe-MBONs induced approach: MBON-g1/g2 (N_exp_=6, N_larvae_≈210) and MBON-m1 (N_exp_=9, N_larvae_≈450). **g.** Activating the MBONs in the tip of vertical lobe had different effects: MBON-e2 (N_exp_=5, N_larvae_≈140) induced a mild avoidance response, whereas activating MBON-e1 (using a *GAL4* line combined to *tsh-GAL80* to eliminate expression in the nerve cord) induced an approach-like response. Note that most avoidance-promoting MBONs are downstream of appetitive-memory compartments (3/5), and the remainder are downstream of compartment with unknown roles in memory formation. By contrast, most approach-promoting MBONs (4/6) are downstream of an aversive-memory compartment, and the remainder are downstream of compartments with unknown roles in memory formation. **h.** Schematic shows two main mechanisms potentially enabling the MB network to switch from encoding positive or neutral to negative valence after aversive learning: i) In MB compartments receiving projections of aversive DANs (*in red*) and where aversive memory can be formed (*e.g.* “Compartment 1”), a synaptic depression (postulated from *Drosophila* adult, Hige et al., 2015b) takes place between the conditioned odor-KCs and approach-promoting MBONs, skewing the balance towards avoidance-pomoting MBONs. ii) A decreased odor-evoked response in inhibitory avoidance-promoting MBONs can disinhibit the avoidance-promoting MBONs downstream of the MB compartments receiving signal of appetitive US (*in blue*) and where appetitive memory can be formed (*e.g.* “Compartment 2”). **i.** Circuit diagram obtained from EM reconstruction (Eichler *et al.*, 2017), overlaid with neurotransmitter profile (Eichler *et al.*, 2017) and behavioral phenotypes (**a-f**), displays lateral inhibition between MBONs of opposite valence. Blue rim = approach-promoting, red rim = avoidance-promoting, grey rim = no behavioral phenotype observed, purple: not tested. Arrows indicate excitatory cholinergic connections, bars indicate inhibitory GABAergic connections, and squares indicate glutamatergic connection, also likely inhibitory in *Drosophila* (Liu and Wilson, 2013), and circles indicate unknown neurotransmitter. Vertical and horizontal bars within neuron nodes indicate source (*i.e.* emitting projections), sink (i*.e*. receiving projections), or transfer MBONs (*i.e*. emitting and receiving projections) within the MBON network.

Activation of some MBONs repressed turning and promoted crawling, compared to controls. Some of these MBONs also promoted turning and repressed crawling in response to a decrease in their activity, at the offset of optogenetic activation. We classify these MBONs as promoting odor approach. If these neurons were activated by an odor (as would occur when the animal is crawling towards an odor source) they would repress turning allowing the animal to approach the odor. If the activity of these neurons is decreased (as would occur if the animal is crawling away from an odor course) they would promote turning.

Activation of other MBONs promoted turning and repressed crawling, compared to controls (Fig. 2b-g). We classify these MBONs as promoting avoidance. If these neurons were activated by an odor (as would occur when the animal is crawling towards an odor source) they would promote turning which would result in odor avoidance.

We found that most MBONs that promote approach innervate compartments implicated in aversive memory formation and receive synaptic input from DANs whose activation (paired with odor) induces aversive memory (Fig. 2e-f and 2h, (Eschbach et al., 2020)). Conversely, most MBONs that promote avoidance innervate compartments implicated in appetitive memory formation and receive synaptic input from DANs whose activation (paired with odor) induces appetitive memory (Fig. 2d and 2h, (Eschbach et al., 2020; Rohwedder et al., 2016; Saumweber et al., 2011). Curiously, some MB compartments of unknown function (Upper Vertical Lobe and Calyx) are innervated by two distinct MBONs that had opposite effects on behavior (Fig. 2b and 2g).

Overall our findings are consistent with mechanisms described in the adult *Drosophila* (Yoshinori Aso, Hattori, et al., 2014; David Owald & Waddell, 2015): the formation of an aversive olfactory memory reduces the conditioned odor drive to approach-promoting MBONs, and vice versa, for appetitive memory.

### Lateral inhibition between MBONs that promote opposite behaviours

In order to begin to understand how the activity of the entire population of MBONs is used to control learnt odor approach or avoidance we first analysed direct interactions between MBONs that promote opposite behaviors. We have recently mapped the synaptic-resolution connectivity between all MBONs in a first instar larval brain and identified their neurotransmitter expression (Eichler et al., 2017). We therefore combined the behavioral effects of MBON activation with this information (Fig. 2i). We observed extensive lateral inhibition between MBONs that promote opposite behaviours: 6 instances of an avoidance-promoting MBON projecting inhibitory synapses (*i.e.* GABAergic or glutamatergic, (W. W. Liu & Wilson, 2013)) onto an approach-promoting MBON, and 9 instances of an approach-promoting MBON projecting inhibitory synapses onto an avoidance-promoting MBON.

We postulate that for these neurons a mechanism of disinhibition could enhance the contrast in odor drive to approach- and avoidance-promoting MBONs (Fig. 2h). In the example of an aversive olfactory memory, the depressed odor drive onto approach-promoting MBONs would be accompanied by reduced inhibition of avoidance-promoting MBONs. Such disinhibition likely takes place following aversive learning in the adult (D. Owald et al., 2015). In the larva, we find that several approach- and avoidance-promoting MBONs are targets of lateral inhibition by MBONs of opposite valence, suggesting that this may be a general principle of MB organisation (Fig. 2h).

### Comprehensive EM reconstruction of all neurons downstream of all MBONs reveals candidate neurons for comparing odor drive to distinct MBONs

To investigate how the circuits downstream of MBONs could compare the odor drive to approach- and avoidance-promoting MBONs to read out the learnt odor valence, we reconstructed all postsynaptic neurons of all 24 MBONs in both the right and left brain hemispheres (Fig. 3a-b, Extended Data Fig. 3, Supplementary Adjacency Matrix). We identified 167 left and right homologous pairs of neurons that were strongly and reliably connected to MBONs (see Methods for definition of strong and reliable, Fig. 3a). We named these neurons MB second-order output neurons (MB2ONs).

**Figure 3:**
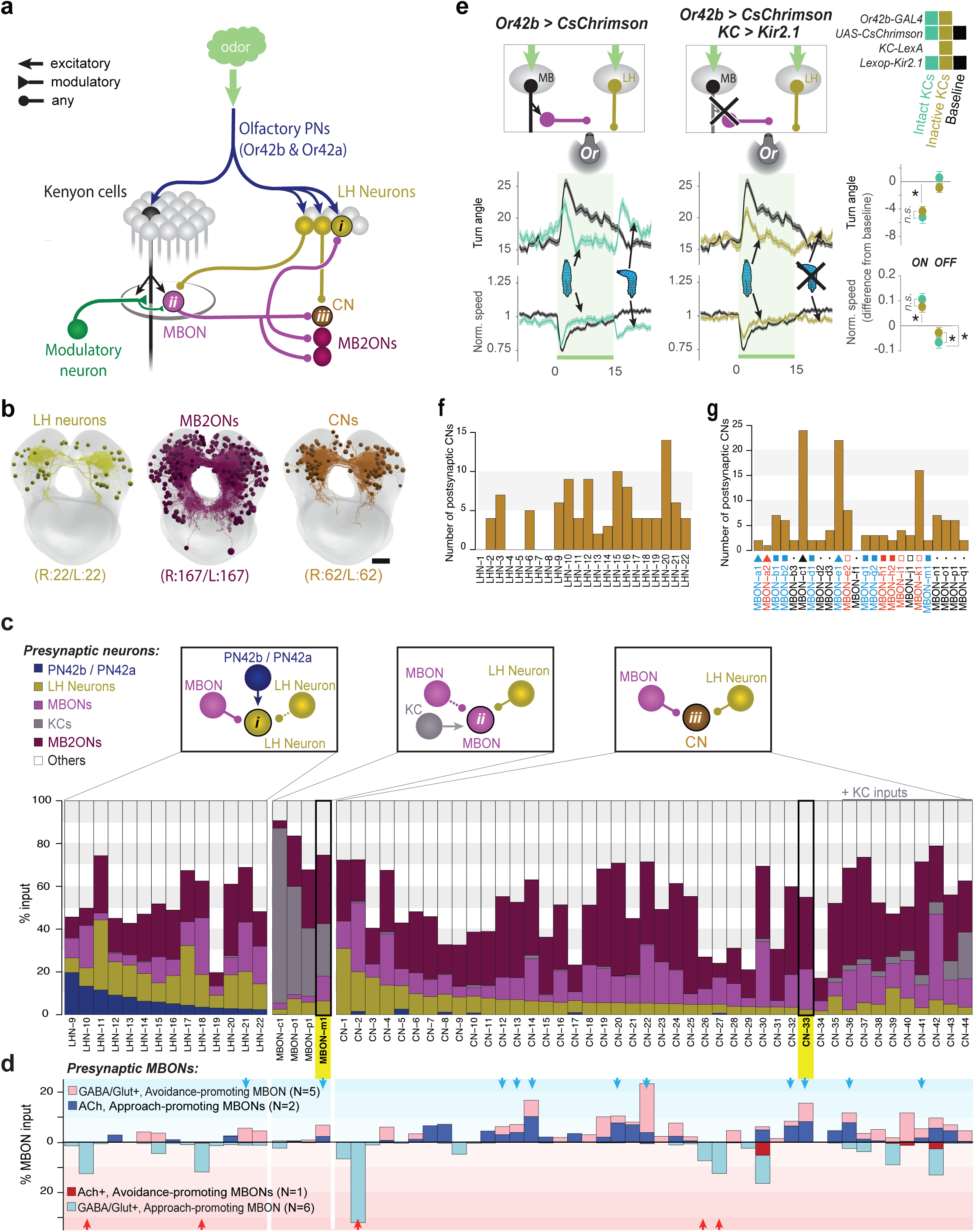
Parallel olfactory pathways interact at different levels. **a.** Schematic of interaction between the LH and MB olfactory pathways. All 167 pairs of neurons postsynaptic to MBONs, so-called MB2ONs (*dark purple*), have been reconstructed from an electron microscopy volume, as well as the 22 pairs of neurons downstream of two olfactory PNs in the LH (*dark yellow*, Extended Data Figure 4). This unraveled three types of convergence in olfactory inputs: i) a direct projection of MBONs onto LH neurons, ii) a direct projection of LH neurons onto MBONs, and iii) common postsynaptic target of LH neurons and MBONs (*i.e.* onto a MB2ON). **b.** Projections of the reconstructed brain neurons. **c.** All neurons receiving convergent inputs from the parallel olfactory pathways LH and MB were called ‘CNs’ for Convergence Neurons. Top schematics show the three different types of convergent interactions described from EM reconstruction in (**a**). The bar graphs show the fraction of inputs received by each CN from different types of neurons: olfactory PNs (Projection Neurons), LH neurons (Lateral Horn), MBONs (Mushroom Body Output Neurons), MB2ONs (2nd Order MBONs), KCs (Kenyon Cells), and other. Shared postsynaptic MB2ONs (type ii) are the most common type of interaction between LH and MB olfactory pathways. Note that some type iii CNs receive direct KC input, but were not classified as MBONs for this study because connections from individual KCs were weak (<3 synapses) or asymmetric (≥3 KC connection(s) only present in one hemilateral partner) (CN-35, -36, -37, -38, -39, -40, -42), KC inputs were axo-axonic (CN-43), or do not target discrete MB compartments (CN-44, Eichler *et al*., 2017). Additionally, none of these type iii CNs received above threshold MBIN input. Also note that some type iii CNs receive asymmetric or subthreshold PN input (*e.g.* CN-5) and are therefore not considered LHNs. **d.** Inputs from MBON are further separated according to their known behavioral effects and neurotransmitters (Fig. 2g and Eichler *et al*., 2017). Some CNs receive excitatory inputs from approach-promoting MBON(s) and inhibitory inputs from avoidance-promoting MBON(s), or inhibitory inputs from multiple avoidance-promoting MBONs (*blue arrows*). A few other CNs integrate inhibitory inputs from multiple approach-promoting MBONs (*red arrows*). This suggests that a valence signal coming from the MBONs is encoded by these CNs. Only neurons with at least 5% of their total input from the appropriate MBONs were considered (*blue arrows*, *red arrows*). **e.** The LH path can drive odor approach. Optogenetic activation of Or42b, an olfactory neuron encoding notably ethyl acetate (Kreher et al., 2008), induced a reduction (compared to the control reaction to light, *black*, N≈350) of turn behavior in naive larvae at light onset and an increase of turn at light offset (*cyan*, N≈250), characteristics of an approach-like response (see Fig. 2a). Silencing KC with Kir2.1 (*dark yellow*, N≈300) did not alter the onset response but abolished the offset response component. Plots on the right show normalized parameters as in Fig. 2b-f. **f.** Distribution of counts of strongly connected postsynaptic CNs downstream of each LHN. Some LH neurons of the Or42a/Or42b pathway target more CNs than others. **g.** Distribution of counts of strongly connected postsynaptic CNs downstream of each MBON. Some MBONs target more CNs than others.

40/167 MB2ONs synapse directly onto MB modulatory neurons, and had been previously reconstructed as part of our investigation of modulatory neuron inputs (Eschbach et al., 2020). We have previously named these neurons, feedback neurons (FBNs). Another 58/167 neurons provide indirect two-step feedback to modulatory neurons by synapsing onto a pre-modulatory neuron (Supplementary Adjacency Matrix, (Eschbach et al., 2020).

We observed both divergence and convergence of MBON inputs at the downstream layer. Many MB2ONs (101/167) receive inputs from only one MBON (Extended Data Fig. 3 and 4a), and each MBON synapses onto multiple MB2ONs (Extended Data Fig. 3 and 4b, Supplementary Adjacency Matrix). Consequently, each MBON projects to a unique combination of MB2ONs (Extended Data Fig. 3 and 4c, Supplementary Adjacency Matrix). Nevertheless, we observed a large population of 66 MB2ON types that received convergent input from multiple MBONs (Extended Data Fig. 3 and 4a). Some integrate input from MBONs that promote the same behaviour (13/66), but many more integrate input from MBONs that promote opposite behaviours (27/66, Extended Data Fig. 3). Interestingly, many of these (18/27) appear to receive excitatory connections from MBONs that promote one behaviour and inhibitory connections from MBONs that promote the opposite behaviour (Extended Data Fig. 3). These MB2ONs could compare the odor drive to MBONs that promote opposite behaviours and thereby compute the learnt valence of an odor based on memory traces from multiple compartments.

### EM reconstruction of LH neurons reveals how LH and MB pathways converge

Next we asked how the learnt valence signals from the MB are integrated with the innate valence signals from the LH, for example, to enable a switch from innate odor approach to learnt avoidance. We therefore sought to 1) identify all the LHNs downstream of olfactory projection neurons (PNs) with strongest response to an innately attractive odor, ethyl acetate, 2) test whether these LHNs support innate odor attraction even in the absence of a functional MB, and 3) determine the patterns of synaptic connections between these LHNs, MBONs, and MB2ONs.

Olfactory receptor neurons (ORNs), ORN42a and ORN42b (Kreher et al., 2005, 2008) show the strongest response to ethyl acetate and synapse onto PN42a and PN42b, which send projections to both the MB and the LH. While all of the olfactory PNs and KCs were recently reconstructed in an EM volume of a first instar larval nervous system (Berck et al., 2016), the neurons downstream of PN42a and PN42b, other than KCs, were previously unknown. We therefore reconstructed all neurons downstream of PN42a and PN42b in the same EM volume and identified 22 pairs of LHNs (LHNs, Fig. 3a-c, Extended Data Fig. 5a-c, Supplementary Adjacency Matrix).

Second, we asked whether the LHNs downstream of PN42b are sufficient to support ORN42b-driven innate odor attraction even in the absence of a functional MB pathway. An increase in the activity of ORN42a or ORN42b in naïve animals decreases turning, while a decrease in ORN42a/b activity increases turning, indicating these neurons promote approach of innately attractive odors (Gepner et al., 2015; Hernandez-Nunez et al., 2015; Schulze et al., 2015). We compared turning in response to optogenetic manipulation of ORN42b activity in larvae with silenced (using the GFP-tagged potassium channel Kir2.1, (Baines et al., 2001) or intact KCs (Fig. 3d). In both groups, we observed a comparable and significant decrease in turning in response to an increase in ORN42b activity (Fig. 3d). This confirms that the LHNs downstream of PN42b can mediate innate odor approach behavior, even in the absence of a functional MB pathway, by repressing turning in response to an increase in odor concentration (as would occur when the animal is crawling towards an attractive odor source (Fig. 3d).

Interestingly, larvae with silenced KCs responded differently to animals with intact KCs to a reduction in ORN42b activity, at the offset of optogenetic activation (Fig. 3d). This suggests that the MB contributes to some aspects of the innate odor response, specifically to increasing turning in response to a decrease in odor concentration (as would occur when the animal is crawling away from an innately attractive odor source). Consistent with this idea, silencing of KC resulted in less efficient navigation in a gradient of ethyl acetate (Extended Data Fig. 6).

Next, we analysed the anatomical patterns of interaction between LHNs, MBONs and MB2ONs. We predicted the signs of connections (inhibitory or excitatory) made by MBONs based on their known neurotransmitter expression (Eichler et al., 2017). Because we have not yet generated GAL4 lines for targeting LHNs in the larva, we could not determine their neurotransmitter identity. Our EM reconstruction revealed direct connections from some MBONs onto 14 LHNs, similar to recent findings in the adult *Drosophila* (Dolan et al., 2018); Fig. 3a,c, Extended Data Fig. 5bi). However, we also observed two new patterns of convergence between LHNs and MBONs. First, direct connections from some LHNs onto four MBONs (Fig. 3a,c, Extended Data Fig. 5bii-iii). Second, the convergence of both LHNs and MBONs onto 44 MB2ONs (Fig. 3c-d, Supplementary Adjacency Matrix). Collectively, we call the neurons that receive LHN and MBON input, “Convergence Neurons” (CNs, Fig. 3c-d, Extended Data Fig. 7a-c). Most LHNs that receive direct MBON input also receive reliable input from other LHNs, and some MBONs (MBON-m1) that receive direct LHN input also receive reliable input from other MBONs, and are therefore also CNs (Fig. 3c). Distinct CNs receive distinct patterns of inputs from LHNs and MBONs (Extended Data Fig. 7a-c, Supplementary Adjacency Matrix). Interestingly, many CNs are also FBNs (18) and synapse directly onto modulatory neurons (we name these CN/FBN, but refer to them as CN here, for brevity (Supplementary Adjacency Matrix, (Eschbach et al., 2020)).

Based on the type of input that the CNs receive from approach- or avoidance-promoting MBONs, we postulated they could fall into at least two different functional classes (Fig. 3d). One class of CNs potentially promotes avoidance, since they receive a significant fraction of input (more than 5%) from inhibitory approach-promoting MBONs (N=5). The second class of CNs potentially promotes approach, since they receive a significant fraction of input from excitatory approach-promoting MBONs and inhibitory avoidance promoting MBONs (N=11). The CNs that integrate inputs of opposite signs from MBONs that promote opposite actions as well as from LHNs potentially compute the overall predicted value associated with approaching or avoiding an odor and promote actions based on these predictions. To test this idea, we asked 1) whether these neurons indeed receive functional inputs from both MB and LH; 2) whether they can drive approach or avoidance.

### MBON-m1 receives functional inputs from LH and MB

Investigating functional connections from LHNs and MBONs onto CNs requires genetic tools for selectively targeting individual CN types. We have not yet generated Split-GAL4 lines for the newly identified CN neuron types, but we have generated Split-GAL4 lines for many MBONs (Extended Data Fig. 1). We therefore focus our initial investigation on MBON-m1 that integrates direct synaptic inputs from LHNs downstream of ORN42b PNs, from KCs, and from other MBONs (Fig. 4a). Specifically, MBON-m1 receives cholinergic (excitatory) input from MBONs that promote approach (MBON-e1) and GABAergic (inhibitory) and glutamatergic (likely inhibitory, (W. W. Liu & Wilson, 2013)) input from MBONs that promote avoidance (MBON-h1, MBON-h2, MBON-i1). KC input is also thought to be cholinergic (excitatory, (Barnstedt et al., 2016)).

**Figure 4:**
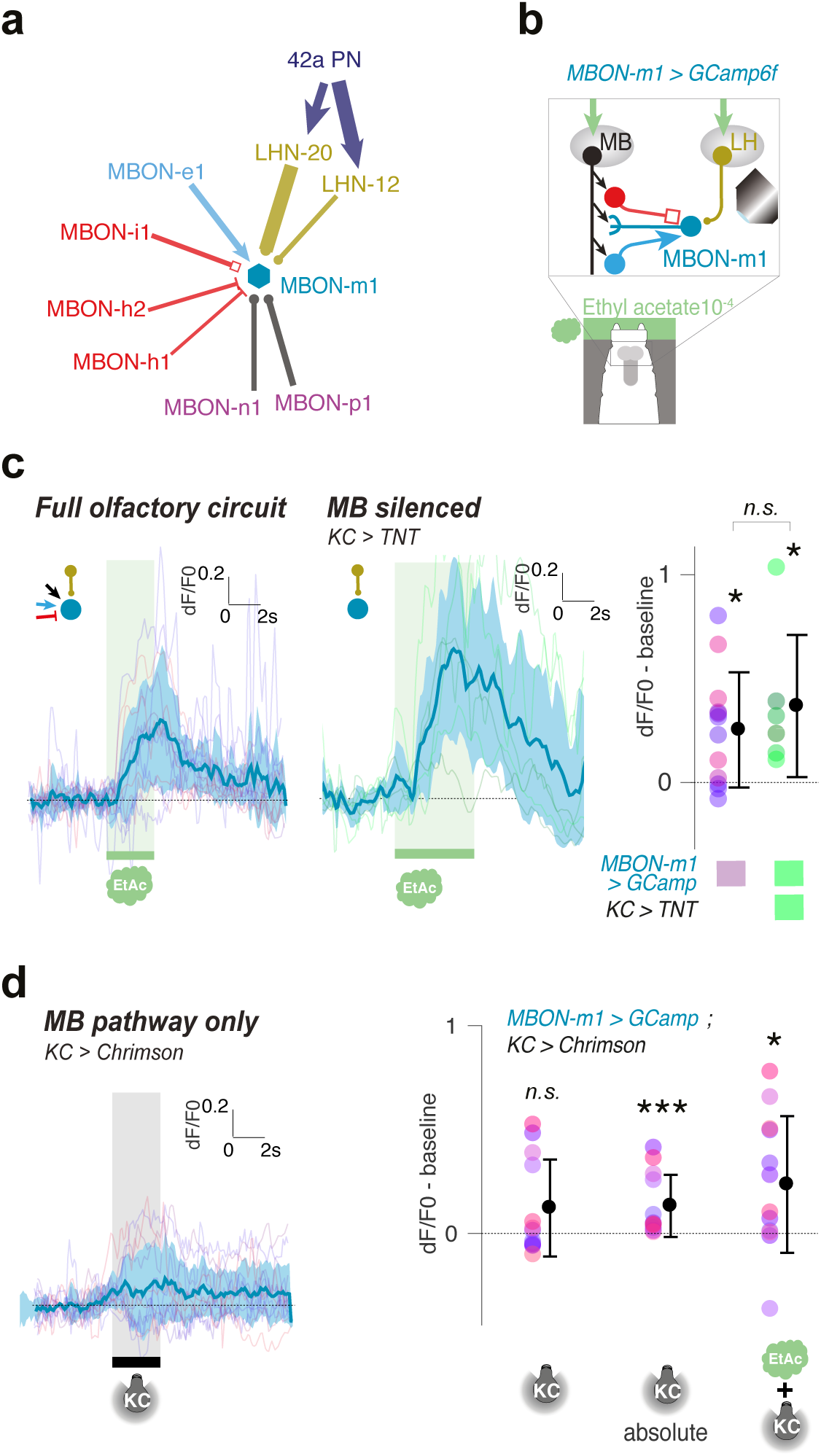
MBON-m1 integrates LH and MB-derived inputs. **a.** EM circuit graph of MBON-m1, a type ii ‘Convergence Neuron’ that integrates inputs from other MBONs and LHNs. In particular, MBON-m1 receives convergent input from the inhibitory avoidance-driving MBON-h1, -h2 (both GABA-positive) and MBON-i1 (VGluT-positive), and from the excitatory approach-driving MBON-e1 (ChAT-positive). **b.** Calcium activity of MBON-m1 was imaged *in vivo* in larvae trapped in microfluidic device (Si *et al.*, 2019) and exposed to the odor ethyl acetate diluted in deionized water (**c**) and/or to optogenetic activation of Chrimson-expressing KCs (**d**). For all plots, curves show fluorescence normalized to baseline (before odor presentation, dF/F0), and scores are calculated for the 3 first seconds of odor presentation. Individual traces are shown in Extended Data Fig. 8. **c.** Regardless of whether the larval olfactory pathways were left intact (left panel, N=12) or the MB pathway was blocked (using TNTe, right panel, N=6), MBON-m1 showed excitatory response to odor exposure. This suggests that, in naïve larvae, MBON-m1 is significantly excited by the attractive odor ethyl acetate mainly via the LH neurons. *: p<0.05, Wilcoxon test. **d.** The average response of MBON-m1 to optogenetic activation of KCs is zero. However, both individual traces and significant normalized response to KC activation in absolute value suggest that the MB pathway can drive MBON-m1 towards either excitation or inhibition. N=12, *: p>0.05, ***: p<0.0001, Wilcoxon test.

To test whether the connections from the LHNs downstream of PN42a/42b were functional and to determine their sign, we compared MBON-m1 calcium responses to ethyl acetate in animals with silenced KCs (by expressing tetanus toxin light chain with GMR14H06-LexA > LexAop-TNTe, (Sweeney et al., 1995)) and controls with functional KCs (Fig. 4b). We verified that the MB pathway is silenced by this method by observing no odor memory after an odor-sugar training protocol (See Methods, Extended Data Fig. 8). We imaged MBON-m1 activity in intact living animals immobilised in a microfluidic device (for improved image quality we used first rather than third instar larvae, (Si et al., 2019)). We found that MBON-m1 was indeed activated by ethyl-acetate both in the presence and absence of a functional MB pathway (Fig. 4b, Extended Data Fig. 9a-b), indicating the LH neurons provide functional excitatory input to MBON-m1. Furthermore, we did not observe a significant difference in calcium response of MBON-m1 in the two conditions, suggesting that the net contribution of the MB to MBON-m1 response to an odor in naive animals may be 0 (Fig. 4c).

To investigate the net contribution of the MB to MBON-m1 activity we also directly optogenetically activated all KCs (using GMR14H06-LexA line to drive CsChrimson) and imaged calcium transients in MBON-m1 (Fig. 4c). To be as close as possible to a naive state we did this in individuals that had never been exposed to specific associative olfactory training. Optogenetic activation of KCs evoked inhibitory responses in some individuals, excitatory responses in some, or no response in others (Fig. 4c, Extended Data Fig. 9c-d). The variability in response of MBON-m1 to KC activation across individuals could be due to different experiences prior to these experiments. On average, across individuals, the net response of MBON-m1 to KC activation is not significantly different from 0 (Fig. 4c). This result further confirms our finding above, that the response of MBON-m1 to natural odor comes largely from the LH pathway in the naive state.

Furthermore a natural odor is expected to activate only a small fraction (ca. 5%, (Honegger et al., 2011)) of KCs. Thus, in our experiments with direct optogenetic activation of all KCs, MBON-m1 likely received a much stronger excitatory input from KCs than in response to a natural odor. Despite this, KC activation did not activate MBON-m1 in many individuals and on average the net response was not significant. MBON-m1 receives a significant fraction of input from inhibitory avoidance-promoting MBONs which likely counterbalance the excitation by KCs and by excitatory MBONs. Consistent with this idea, the absolute value of the response of MBON-m1 to KC activation was significant, indicating that there is a response, which is sometimes excitatory and sometimes inhibitory (Fig. 4c).

Together these experiments provide support for the prediction from the connectome that MBON-m1 integrates functionally excitatory inputs from an approach-promoting LH pathway with functionally excitatory (from KCs and from approach-promoting MBONs) and inhibitory (from avoidance-promoting MBONs) input from the MB.

### MBON-m1 bi-directionally controls turning and contributes to odor approach

Since MBON-m1 receives functional input both from the LH and the MB, it could contribute to innate olfactory behavior, but its activity could also be modified by learning. First, we wanted to explore in more detail the role of MBON-m1 in behaviour. We found that optogenetic activation of MBON-m1 represses turning (Fig. 2f). In contrast, we observed the opposite response at the offset of optogenetic activation (Fig. 2f), suggesting that a decrease in MBON-m1 activity relative to the baseline may promote turning. To confirm this we acutely optogenetically hyperpolarized MBON-m1 with the anion channel-rhodopsin GtACR2 (Govorunova et al., 2015; Mohammad et al., 2017) and found this increased turning (Fig. 5a). Thus, increasing and decreasing MBON-m1 activity relative to the baseline, was sufficient to decrease and increase turning, respectively. This suggests MBON-m1 plays a role in approaching an attractive odor source, by increasing and decreasing turning in response to a decrease and an increase in odor concentration, respectively. To test this we constitutively hyperpolarized MBON-m1 by expressing Kir2.1 in groups of naive larvae and recorded their behavior in a gradient of an innately attractive odor, ethyl acetate (Fig. 5b). Indeed, we found that silencing MBON-m1 resulted in suboptimal chemotaxis in naive animals (Fig. 5b).

**Figure 5:**
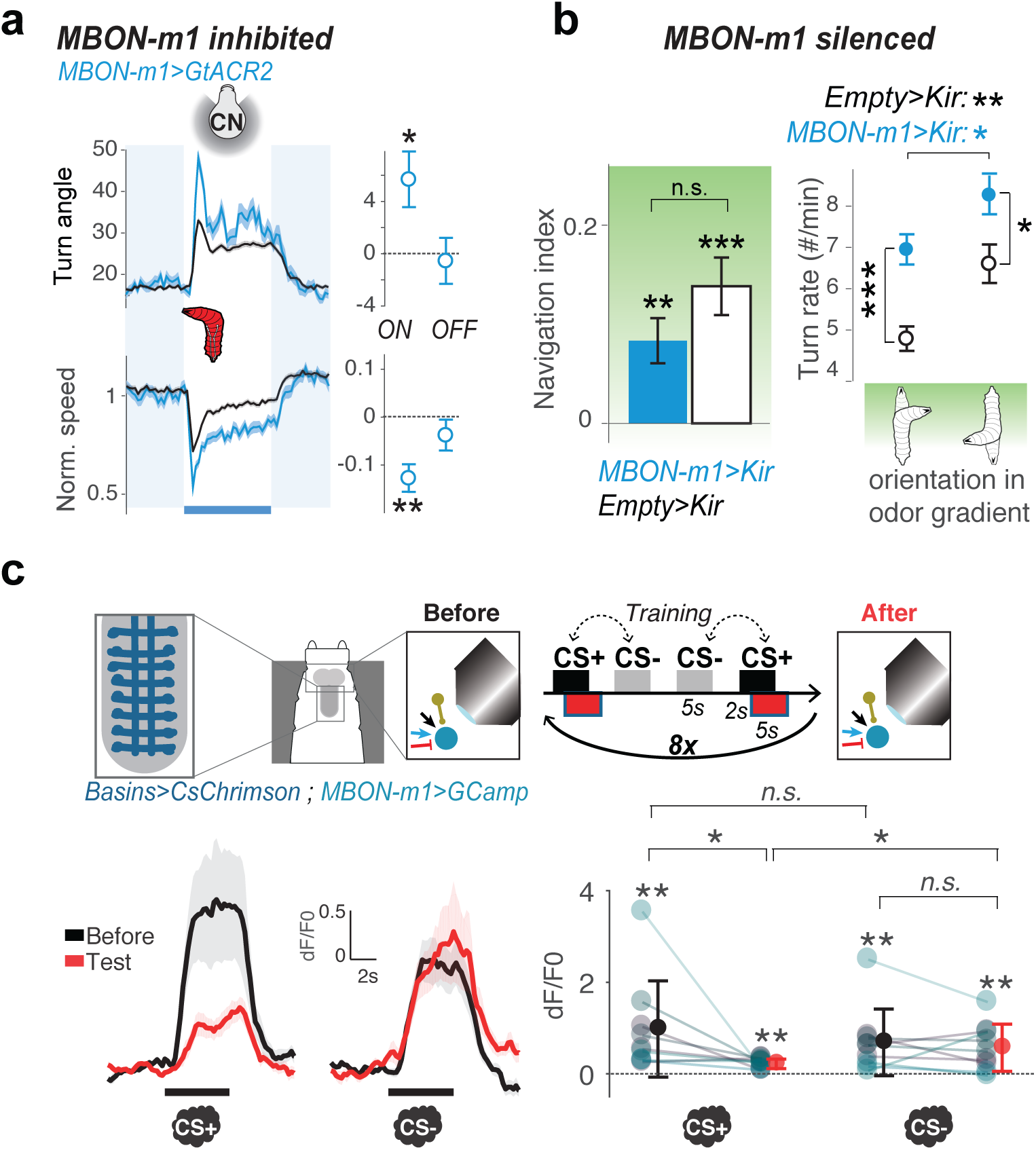
Odor response of MBON-m1 is modified by experience. **a.** Optogenetic activation of MBON-m1 leads to decreased turn at onset and increased turn at offset (Fig. 2f). Fittingly, optogenetic inactivation of MBON-m1 (via the hyperpolarizing channelrhodopsin GtACR2) induces increased turn, similar to the activation offset response. Right plot shows normalized differences in turn angle and speed as in Fig. 2b-g. Thus, MBON-m1 promotes approach when excited, and avoidance when inhibited. *: p < 0.05, **: p<0.001, in Welch’s Z test. **b.** MBON-m1 is involved in navigation behavior. Silencing MBON-m1 with Kir2.1 affects innate approach to the odor source (*left panel*), likely because of a general increase in turn frequency, unspecific to larval orientation (*right panel*). *: p < 0.05, **: p<0.001, ***: p<0.0001 in Welch’s Z test. **c.** Calcium activity of MBON-m1 was imaged *in vivo* in larvae trapped in a microfluidic chip, exposed to odors and to the optogenetic activation of nociceptive neurons (‘Basins’) located in the nerve cord. This fictive punishment was delivered 2s after the exposure to the paired odor (CS+). The response of MBON-m1 to CS+ presentation before and after the pairing is compared to the response to CS- (another odor presented without punishment). The response to CS+, but not to CS-, significantly decreased after training (N=9). Curves show average fluorescence normalized to baseline (*i.e.* before odor presentation) +/- s.e.m. Plots show mean fluorescence scores during odor delivery per animal, and their average +/- s.e.m. in black or red. The odors presented were different for different animals and were one of the following 4 combinations of CS+/CS- odors: AM+/EA- (N=4), EA+/AM- (N=3), EA+/ME- (N=1), ME+/EA- (N=1). *: p<0.05, **: p<0.005 in a Wilcoxon test for paired comparison.

### Aversive learning depresses MBON-m1 response to conditioned odors

We also wanted to confirm that MBON-m1 conditioned-odor response is modified by learning. MBON-m1 receives direct input from DANs that are activated by aversive stimuli and whose activation paired with odor induces aversive memory (Eschbach et al., 2020). Based on studies in adult *Drosophila* (Hige, Aso, Modi, et al., 2015; Hige, Aso, Rubin, et al., 2015; Sejourne et al., 2011) we predicted that aversive learning would reduce the conditioned odor drive to the approach-promoting MBON-m1. To test this we asked whether pairing an innately attractive odor with punishment (optogenetic activation of Basin interneurons, (Ohyama et al., 2015) depresses the response of MBON-m1 to that odor. We performed these experiments in first instar larvae immobilized in a microfluidic device (Fig.5c, Extended Data Fig. 9e, 10a-b, (Si et al., 2019)). As expected, we found significantly decreased responses of MBON-m1 to the paired (CS+) and not the unpaired odor (CS-, Fig. 5c). Aversive learning could reduce the conditioned odor drive to MBON-m1 in two ways: i) through depression of the KC-to-MBON-m1 connections due to the pairing of KC activation with DAN-g1, -g2, and -d1 activation (activated by punishment, (Eschbach et al., 2020)); ii) aversive learning in other compartments could disinhibit MBONs that inhibit MBON-m1 (*i.e*. reduced activation of approach-promoting MBON-g1 and -g2 would disinhibit avoidance-promoting MBON-i1). Together our findings are consistent with the idea that MBON-m1 is activated by innately attractive odors (via the LH pathway) and promotes approach and that aversive learning reduces approach by depressing the conditioned odor drive onto MBON-m1.

### CN-33 receives functional inputs from LH and MB

MBON-m1 is just one of a population of 11 CNs predicted based on connectivity to be activated by approach-promoting LHNs and MBONs and inhibited by avoidance promoting MBONs (Fig. 3c). The other 10 do not receive significant KC input so learning must modulate their activity indirectly, via MBONs. We wanted to functionally investigate at least one other member of this class. We had a Split-GAL4 line (Pfeiffer et al., 2010), SS02108 (Eschbach et al., 2020), that drives gene expression in a CN that we called CN-33/FAN-7. This neuron was previously shown to provide feedback to DANs, although its LH inputs were unknown (Eschbach et al., 2020). Like MBON-m1, it receives cholinergic input from MBONs whose activation promotes approach (MBON-e1) and glutamatergic (likely inhibitory) input from MBONs whose activation promotes avoidance (MBON-i1 and MBON-e2). It also receives input from the same two LHNs downstream of PN42a/PN42b as MBON-m1 (Fig. 6a).

**Figure 6:**
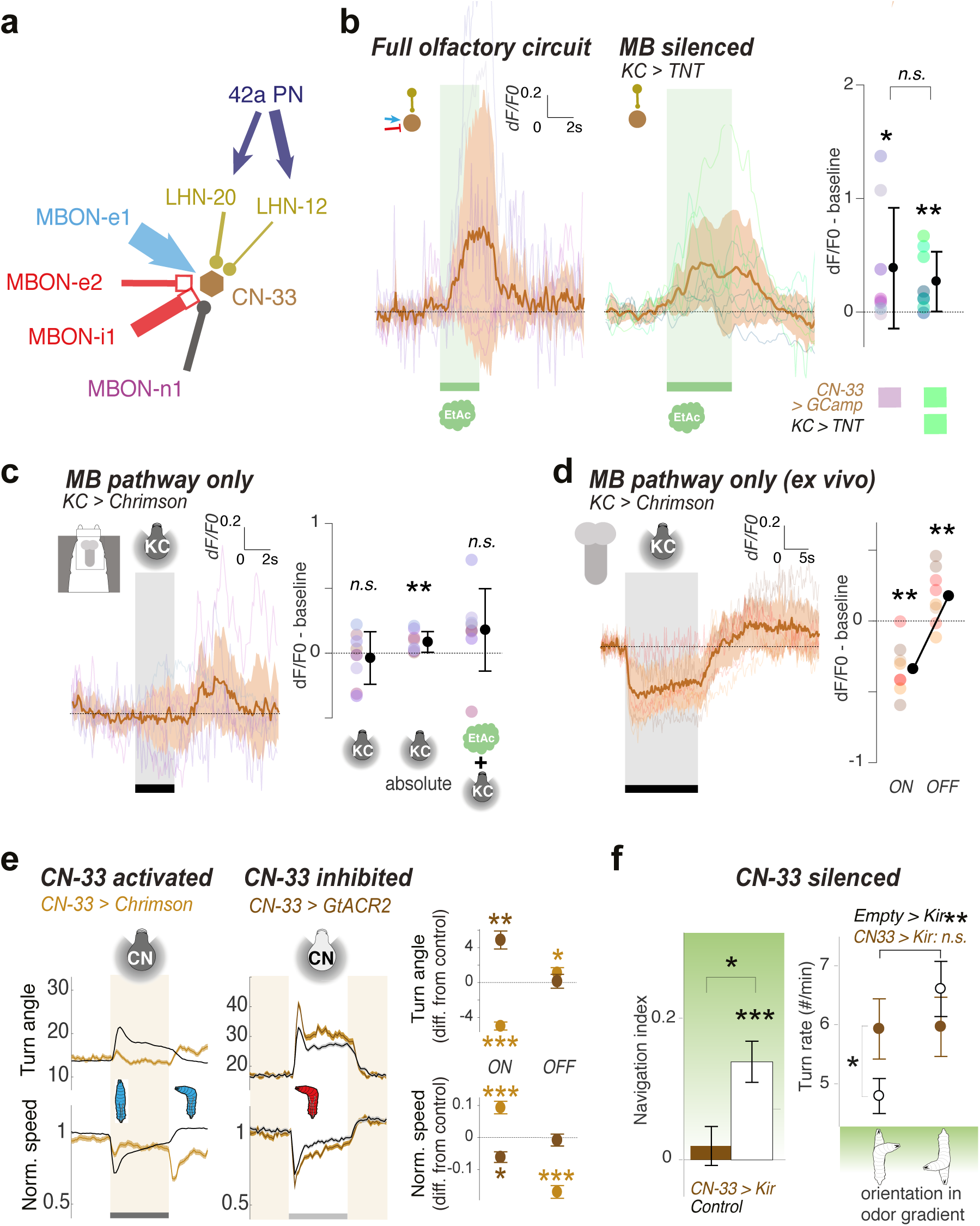
CN-33 integrates LH and MB-derived inputs. **a.** EM circuit graph of CN-33, a ‘Convergence Neuron’ that receives direct input from different MBONs and some LH neurons. In particular, it receives convergent input from the inhibitory avoidance-driving MBON-e2 and MBON-i1, and from the excitatory approach-driving MBON-e1. **b.** Calcium activity of CN-33 was imaged *in vivo* in larvae either with olfactory pathway left intact or with MB pathway blocked. An excitatory response to the odor was observed in CN-33 when both MB and LH pathways were intact. N=8, *: p=0.016, Wilcoxon test. An excitatory odor response was also observed when MB was blocked using TNTe, suggesting excitatory drive from the LH neurons. N=8, *: p=0.016, Wilcoxon test. In all plots, curves show fluorescence normalized to baseline (before odor presentation, dF/F0), and scores are calculated for the 3 first seconds of odor presentation. Individual traces are shown in Extended Data Fig. 10. To figure how the MB pathways shape the odor response in CN-33, we directly activated the Kenyon Cells optogenetically in brain explant (**c**) as well as *in vivo* (**d**). **c.** When CN-33 was imaged *in vivo*, on average, CN-33 did not show a response to KC activation, but showed a slight excitation at light offset, possibly indicating inhibitory rebound. The observation of individual traces however suggests overall MB output neurons drive may be either excitatory or inhibitory, consistent with the description of the connectivity. N=8, Wilcoxon test: p>0.05. **d.** Activation of KCs in brain explants consistently induced an inhibitory response at light onset and an excitatory response at light offset. This suggests that MB output neurons mostly inhibit CN-33. N=8, **: p=0.008, Wilcoxon test comparison to before stimulation. **e.** Optogenetic activation of CN-33 leads to decreased turn at onset and increased turn at offset. Optogenetic inactivation of CN-33 induces increased turn, similar to the activation offset response. Right plot shows normalized differences in turn angle and speed as in Fig. 2b-g. Thus, CN-33 promotes approach when excited, and avoidance when inhibited (See Extended Fig. 8 for more control experiments). *: p < 0.05, **: p<0.001, ***: p<0.0001 in Welch’s Z test. **f.** CN-33 is required for efficient navigation in a gradient of ethyl acetate. When CN-33 was silenced with Kir2.1, the larvae did not approach the odor source (left panel), likely due to the fact that they did not modulate turn frequency when facing or being away from the gradient (right panel, see also Extended Fig. 8). *: p < 0.05, **: p<0.001, ***: p<0.0001 in Welch’s Z test.

To functionally test the contributions of the LH and MB to CN-33 activity, we performed the same kind of imaging experiments as we did for MBON-m1 (Fig 6b-d). To test whether the connections from the LH neurons were functional, we compared CN-33 calcium responses to ethyl acetate in animals with silenced KCs and controls with functional KCs (Fig. 6b, Extended Data Fig. 11a-b). We imaged CN-33 activity in intact animals immobilised in a microfluidics device. We found that CN-33 was indeed activated by ethyl-acetate both in the presence and absence of a functional MB pathway (Fig. 6b), indicating the LH neurons provide functional excitatory input to CN-33.

Next, we wanted to investigate the MB drive onto CN-33 in naive animals. Since, CN-33 receives excitatory inputs from some (approach-promoting) MBONs and potentially inhibitory (glutamatergic) input from other (avoidance-promoting) MBONs, we envisaged three possible scenarios: i) the excitation and inhibition are balanced, consistent with the current working model (David Owald & Waddell, 2015); ii) the inhibition is stronger than excitation with MB potentially reducing the innate response of CN-33 to odor; iii) the excitation is stronger with MB potentially facilitating the innate response of CN-33 to odor. We optogenetically activated all KCs in individuals that had never been exposed to specific associative olfactory training and imaged calcium transients in CN-33, either in intact living animals immobilised in the microfluidics device (Fig. 6c, Extended Data Fig. 11c-d), or in extracted central nervous systems (Fig. 6d, Extended Data Fig. 11e). Optogenetic activation of KCs in intact animals evoked inhibitory responses in some individuals, excitatory responses in some, or no response in others (Fig. 6c). On average, across individuals, there was no significant response to the onset of KC activation, suggesting excitation from approach-promoting MBONs and inhibition from avoidance-promoting MBONs are balanced under these conditions. The variability in response of CN-33 to KC activation across individuals could be due to different experiences across individuals prior to these experiments. Interestingly, in the extracted nervous system, KC activation reliably reduced calcium signals in CN-33, relative to the baseline, indicating that the inhibitory MB drive is stronger than the excitatory one under these conditions (Fig. 6d). This could potentially be explained by the formation of an intense generalised aversive olfactory associative memory during dissection. Alternatively, the relative strengths of excitatory and inhibitory connections in the network could be influenced by state (which could differ between dissected and intact individuals and across individuals). Regardless, the experiments described in this section show that CN-33 is activated by innately attractive odors via the LH pathway, and that it is excited by some (approach-promoting) and inhibited by other (avoidance-promoting) MBONs.

### CN-33 bi-directionally controls turning and contributes to odor approach

Since CN-33 is activated by an innately attractive odor via the LH pathway, we hypothesized that its activation promotes approach. To test this we asked whether optogenetically increasing the activity of CN-33 would repress turning (using SS02108 > UAS-CsChrimson). SS02108 drives expression in two neurons with similar morphology in each hemisphere, CN-33 and MB2ON-86, and weakly in segmentally repeated local interneurons in the nerve cord downstream of aversive mechanosensory neurons. As a control, we removed SS02108-driven nerve cord expression using the Split-GAL4 repressor Killer Zipper (Dolan et al., 2017) under the control of teashirt-LexA (J.-M. Knapp and J. Simpson, unpublished data) promoter (Extended Data Fig. 12a-b). As a second control, we also activated MB2ON-86 alone using SS04330 (see (Eschbach et al., 2020), Extended Data Fig. 12c-d). We found that optogenetic activation of CN-33 decreased turning and increased crawling (Fig. 6e, Extended Data Fig. 12a-d). Additionally, we observed increased turning at the activation offset. To confirm this another way we acutely optogenetically hyperpolarized CN-33 with GtACR2 (Govorunova et al., 2015; Mohammad et al., 2017) and found this increased turning relative to controls (Fig. 6e). Thus, increasing and decreasing CN-33 activity relative to baseline, decreased and increased turning, respectively. This finding is consistent with the idea that CN-33 contributes to odor approach by 1) repressing turning when it is activated by an increase in concentration of an attractive odor; 2) promoting turning when it is inhibited by a decrease in concentration of an attractive odor.

### Silencing CN-33 impairs approach of innately attractive odors

The observed pattern of MBON connectivity onto CN-33 suggests that associative learning can bidirectionally modify CN-33 responses to conditioned odor. Appetitive learning would depress conditioned odor drive to the inhibitory (MBON-i1). This would increase excitation of CN-33 by the conditioned odor thereby increasing approach. Aversive learning would disinhibit the inhibitory (MBON-i1). This would increase inhibition of CN-33 by the conditioned odor thereby decreasing approach or even promoting avoidance.

To test this idea we asked whether inhibiting CN-33 activity reduces approach of an innately attractive odor. We constitutively hyperpolarized CN-33 by expressing Kir2.1 in groups of naive larvae using *SS02108* and recorded their behavior in a gradient of ethyl acetate (Fig. 6f). We found that attraction to ethyl acetate in these manipulated larvae was almost abolished (Fig 6f, see Extended Data Fig. 12e for control line *SS04330*).

Altogether, our findings are consistent with the idea that an increase in CN-33 activity signals an increase in the concentration of an attractive odor and promotes approach (by repressing turning and promoting crawling), whereas reducing the activation of CN-33 reduces approach.

## Discussion

Selecting whether to approach or avoid specific cues in the environment is essential for survival across the animal kingdom. Many cues have both innate values acquired through evolution and learnt values acquired through experience that can guide action selection (Fig. 1). Innate and learnt values are thought to be computed in distinct brain areas (Choi et al., 2011; Q. Li & Liberles, 2015; Marin et al., 2002; Sosulski et al., 2011), but the circuit mechanisms by which they are integrated and by which learnt values can override innate ones are poorly understood. Using the tractable *Drosophila* larva as a model system, we describe with synaptic resolution the comprehensive pattern of convergence between the output neurons of a learning center (the MB) and an innately attractive pathway in the lateral horn (LH). We discovered a population of convergence neurons (CNs) that integrate direct synaptic input from LHNs and MBONs (Fig. 3). We show some CNs are activated by innately attractive odors via the LH pathway and contribute to innate odor attraction (Fig. 4c, 5b, 6b,f). These CNs also receive direct excitatory and inhibitory input from MBONs that promote approach and avoidance, respectively (Fig. 2, 4a,d, 6a,c,d). Learning can therefore alter the balance of excitation and inhibition onto CNs by modifying the conditioned-odor drive to MBONs (Fig. 5c). Together our study provides insights into the circuit mechanisms that allow learnt values to modify innate ones.

### Patterns of convergence between neurons encoding innate and learnt values

The brain areas that compute innate and learnt values of stimuli have been postulated to interact with each other (Heisenberg, 2003; Root et al., 2014; M. Schleyer et al., 2011; Sosulski et al., 2011; Wystrach et al., 2016), but few examples of such interactions have been described to date (Dolan et al., 2018, 2019; Lerner et al., 2020; Sejourne et al., 2011). In principle, MBONs could synapse directly onto LHNs thereby directly modifying innate values. Alternatively, LHNs could directly synapse onto MBONs. Finally, learnt and innate values could initially be kept separate, and MBONs and LHNs could converge on downstream neurons. We have found some MBONs synapse directly onto some LHNs (Fig. 3a,c,d, Extended Data Fig. 5bi), consistent with previous studies in the adult (Dolan et al., 2018). However, we also identified two novel patterns of convergence. First, we found that some MBONs received direct synaptic input from LHNs (Fig. 3a,c,d, Extended Data Fig. 5bii). Second, we found that many MBONs and LHNs converge onto downstream CNs (Fig. 3a,c,d). One MBON (m1) was also a CN receiving significant direct input from other MBONs and from LHNs (Fig. 3a,c,d, Extended Data Fig. 3). Overall the architecture suggests some early mixing of representations of innate and learnt values, but for the most part these representations are kept separate in the LH and MB, and then integrated by the downstream layer. Maintaining some initial separation of representations of innate and learnt values prior to integration could offer more flexibility and independent regulation, for example by context or internal state.

### Convergence neurons compare the odor drive to approach- and avoidance-promoting MBONs

The prevailing model of MB function in adult *Drosophila* proposes that in naive animals the odor drive to MBONs that promote approach and avoidance is equal such that their outputs cancel each other out (David Owald & Waddell, 2015) and the LH circuits guide olfactory behavior (Heimbeck et al., 2001; Q. Li & Liberles, 2015). Learning alters the overall output towards avoidance or approach-promoting MBONs by modifying specific KCs-to-MBON connections. This model raises several important questions. First, how is the output from approach and avoidance-promoting MBONs integrated? Second, how does it interact with the output of the LH? Our findings provide support for this model and shed insight into these questions.

We found a comparable number of MBONs whose activation promotes approach (reduced turning and increased crawling) or avoidance (increased turning and decreased crawling, Fig. 2). Furthermore, we have identified a population of CNs that integrate input of opposite signs from MBONs that promote opposite actions (Fig. 3d). For two members of this class (MBON-m1 and CN-33) we have shown that their net MB drive is 0 across a population of randomly chosen intact larvae that received no prior training (Fig. 4d, 6c, Extended Data Fig. 9, 11). In some individuals the MB drive onto the CNs was excitatory, in some inhibitory, and in others it was 0, showing that the MB can provide both excitatory and inhibitory drive to the CNs. Together our findings are consistent with a model in which these CNs compare odor drive to approach- and avoidance promoting MBONs which is balanced in naive animals. Associative learning can alter the balance of excitation and inhibition onto these CNs by skewing the conditioned-odor drive towards approach- or avoidance promoting MBONs.

### The learning centre contributes to aspects of the innate behavior

Our findings also suggest there might be some exceptions to the general model presented above and that the MB might contribute to some aspects of innate olfactory behavior. While in intact animals we observed no net MB drive onto CNs, in dissected nervous systems the net drive was inhibitory for CN-33 (Fig. 6d, Extended Data Fig. 11). This result could be explained in two ways: either the dissection induced a generalised aversive memory and skewed the MB drive towards inhibition or the net output could be skewed towards inhibition by context, independently of learning. For example, in adult flies, the MB has been implicated in adjusting innate CO2 avoidance to external or internal context (Bräcker et al., 2013; Lewis et al., 2015).

Furthermore, while the MB may not contribute to the innate response to an increase in odor concentration, our results suggest it may contribute to the response to a decrease in odor concentration. Thus, the LH pathway alone was sufficient to repress turning following an increase in activity of olfactory receptor neurons (this strategy enables crawling towards an innately attractive odor source, Fig. 3e). However, animals with silenced KCs did not increase turning following a decrease in the activity of olfactory receptor neurons (this strategy prevents crawling away from an attractive odor source, Fig. 3e). We also found that animals with silenced KCs were less efficient in navigating an attractive odor gradient than controls (Extended Data Fig. 6). Thus, the net MB output may contribute to aspects of innate odor navigation to improve accuracy or efficiency, and its importance could potentially increase with task difficulty. Notably, a defect in innate odor attraction at low odor concentration has been observed in adult *Drosophila* with silenced KCs (Wang et al., 2003). Task-dependent recruitment of additional high order sensory processing areas is widespread in mammals including humans (*e.g.* (Savic et al., 2000; Winstanley & Floresco, 2016; Yue et al., 2017)).

### Lateral inhibition between neurons of opposite value could enhance action selection

Odors can acquire learnt values that are in opposition with the innate ones. How do the learnt values modify the innate ones? How is conflict between opposite values resolved during action selection? One possibility could be reciprocal inhibition between neurons that promote opposite actions enabling one action drive to override another through a winner-take-all mechanism (*e.g.* (Jovanic et al., 2016; Koyama & Pujala, 2018; Seeds et al., 2014). Another possibility could involve the integration of conflicting signals into a unified representation of predicted value, a notion similar to common currency valuation of options (Levy & Glimcher, 2012; Pearson et al., 2014), which could then be used to compute directional choice (*e.g.* (Paisios et al., 2017; Wystrach et al., 2016).

We did not find reciprocal inhibition, but have found examples of lateral inhibition between neurons that promote opposite actions, both between MBONs (Fig. 2i), and from MBONs to LHNs (Extended Data Fig. 5bii), similar to findings in *Drosophila* adult (Cognigni Felsenberg J. and Waddell S., 2018; Dolan et al., 2018, 2019). Lateral inhibition could facilitate action selection by increasing the contrast in activity between approach-and avoidance promoting neurons. However, there are LHNs and MBONs that promote approach and avoidance that are not inhibited by the opposite group (Extended Data Fig. 5bi-ii). Thus, the observed pattern of lateral inhibition does not appear sufficient to enable one pathway to fully suppress the other.

### CNs compute the overall predicted value and promote actions based on these predictions

Rather than being engaged in reciprocal inhibitory interactions with each other, we found that many neurons that promote opposite actions converge onto CNs (Fig. 3a, c). 11 CNs receive excitatory input from LHNs in the attractive pathway and from MBONs that promote approach and inhibitory input from MBONs that promote avoidance (Fig. 3d). For two of these CNs we confirmed they integrate excitatory LH input with both inhibitory and excitatory MB input (Fig. 4a-d and 6a-d). Across individuals, the net MB drive onto these CNs was variable and ranged from inhibition, to excitation, to 0 (Extended Data Fig. 9c and 11c), suggesting that their net MB drive could be modified by learning (Fig. 5c). We have also shown these CNs bi-directionally promote opposite actions: an increase in their activity promotes approach (decreased turing and increased crawling), and a decrease in their activity relative to the baseline promotes avoidance (increased turning and decreased crawling, Fig. 2f, 5a, 6e). Inhibiting these CNs impaired approach of innately attractive odors (Fig. 5b and 6f). Together our findings suggest that i) these CNs compute the overall predicted value of approaching or avoiding an odor by integrating innate and learnt values and ii) promote actions based on these predictions. An increase in their activity signals positive value, and a decrease signals negative value. Our findings are consistent with the following model of action selection (Fig. 7). CNs whose activation promotes approach are activated by innately attractive odors in naive animals via the LH pathway (when the net MB output onto them is 0, Fig. 4d and 6c). Learning could bidirectionally skew the net MB output onto the CNs towards excitation (appetitive learning), or inhibition (aversive learning). After aversive learning the conditioned odor would activate these CNs less (Fig. 5c) resulting in less approach, or sometimes even inhibit them thereby inducing avoidance. This mechanism could have several advantages. If the same neuron mediates both approach and avoidance, depending on whether its activity increases or decreases, the downstream systems would never receive conflicting commands. In addition, it allows for a graduated signal that can generate a range of strong to weak avoidance or approach behaviors.

**Figure 7:**
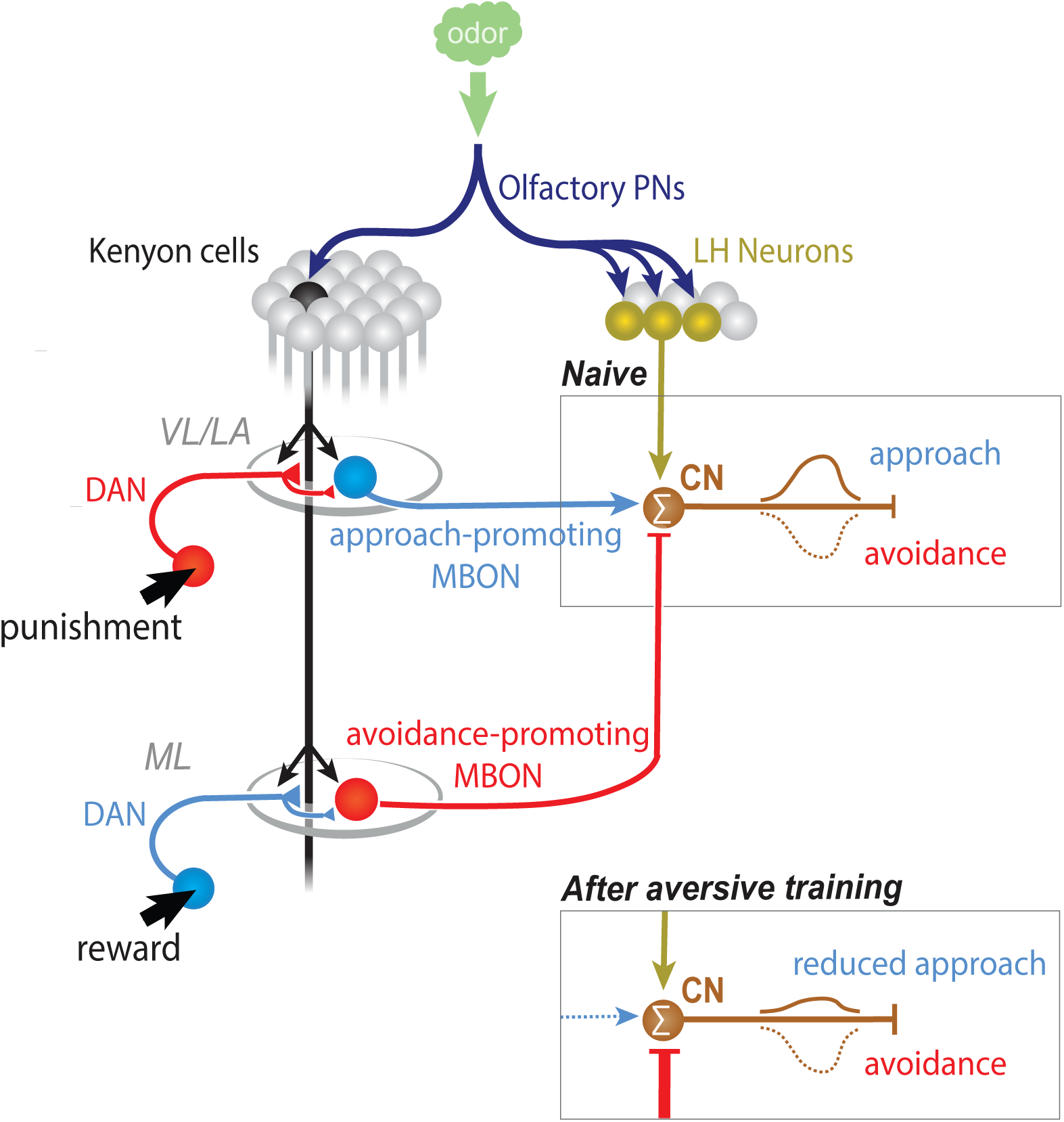
MB and LH pathway both contribute to shaping odor response. Summary diagram of the MB and LH pathways for high-order odor processing and their contribution to odor navigation. In naïve animals, an attractive odor such as ethyl acetate elicits excitatory response in convergence neurons of positive valence such as MBON-m1 or CN-33, mostly via LH neurons, which participate in building the approach to the odor. If an attractive odor is paired with fictive punishment, the response to the paired odor can be reduced in both these neurons, and can shift larval behavior towards odor aversion.

However, an opposite CN population could also exist: those whose activation and inhibition signal negative and positive value (and promote avoidance and approach), respectively (Fig. 3d). CNs whose activation signals positive values could reciprocally inhibit each other. Future comprehensive functional characterisation of all CNs and LHNs identified in this study will reveal this.

### A behavior signal feeds back onto MB modulatory neurons

Interestingly, we find that many of the CNs that receive input from both MBONs and LHNs also provide direct feedback to MB modulatory neurons that provide teaching signals for learning (n=18). In a parallel study (Eschbach et al., 2020) we have shown that at least some of these feedback connections are functional and can influence memory formation. For example, CN-33/FAN-7 is capable of generating an olfactory memory when it is paired with an odor. This type of connectivity is consistent with learning theories that propose that future learning is influenced by predictions based on prior learning (Schultz, 2015). A major role of the CNs discovered in this study may therefore be not only to organize current actions, but also to regulate learning in order to improve future actions based on prior experience. The connectome of the circuits downstream of the learning centre output neurons presented here offers the potential to comprehensively elucidate the neural mechanisms responsible for integrating phylogenetic and individual life history into behavior.

## Methods

### Larvae

Larvae were reared in the dark at 25°C in food vials. The food was supplemented with trans-retinal (SIGMA R2500) at a final concentration of 500μM if the genome contained the *UAS-CsChrimson* transgene. For behavior experiments, the larvae were selected at their third-instar stage; for imaging experiments, they were at first-instar. We verified that first-instar larvae were capable of learning using optogenetic manipulations (Extended Fig. 5).

To train larvae with optogenetic punishment, we used the *GMR72F11-GAL4* (Ohyama et al., 2015); BDSC 39786) line crossed to *20XUAS-CsChrimson-mVenus* (Klapoetke et al., 2014); BDSC 55134). To assess the response to optogenetic activation of MBONs or CNs, Split-GAL4 (Pfeiffer et al., 2010) were crossed to direct the expression of *20XUAS-CsChrimson-mVenus*. The empty stock *y w;attP40;attP2* (Pfeiffer et al., 2010) was crossed to the effector as a baseline control. The following driver lines were used:

| Neuron | SplitGAL4 | AD | DBD | Figure | Reference |
| --- | --- | --- | --- | --- | --- |
| MBON-a1 | *SS02006* | *93G12* | *71E06* | 2 | unpublished |
| MBON-a2 | *SS01417* | *52E12* | *93G12* | 2 | unpublished |
| MBON-b1 + MBON-b2 | *SS01708* | *12G03* | *21D02* | 2 | unpublished |
| MBON-c1 | *SS01776* | *74B11* | *20F01* | 2 | unpublished |
| MBON-d1 | *SS01705* | *11E07* | *52H01* | 2 | unpublished |
| MBON-d2 | *SS04231* | *121A04* | *87G02* | 2 | unpublished |
| MBON-d3 | *SS24027* | *111B05* | *100H11* | 2 | unpublished |
| MBON-e2 | *SS04559* | *65A05* | *102D01* | 2 | unpublished |
| MBON-g1 or g2 | *SS02130* | *23B09* | *21D06* | 2 | unpublished |
| MBON-h1 or h2 | *SS00894* | *67B01* | *11F03* | 2 | unpublished |
| MBON-i1 | *SS01726* | *20C05* | *14C08* | 2 | (Eschbach et al., 2020) |
| MBON-j1 | *SS01972* | *128F10* | *12C11* | 2 | (Eschbach et al., 2020) |
| MBON-k1 | *SS01962* | *VT033301* | *27G01* | 2 | (Saumweber et al., 2018) |
| MBON-m1 | *SS02163* | *52H01* | *40F09* | 2 and 6 | (Eschbach et al., 2020) |
| CN-4 + MB2ON-86 | *SS02108* | *13D05* | *40F09* | 4 and 5 | (Eschbach et al., 2020) |
| MB2ON-86 | *SS04330* | *17H07* | *40F09* | Ext. data | (Eschbach et al., 2020) |

For a few MBONs, GAL4 lines inserted at the *attP2* site (from the FlyLight GAL4 collection, (Jenett et al., 2012) were crossed to *UAS-CsChrimson; tsh-GAL80*, to antagonize effector expression in the ventral nerve cord. *Tsh-LexA* was a gift from J.-M. Knapp and J. Simpson (unpublished stock). In brief, the *tsh-LexA* driver is an enhancer trap inserted into the 5′ UTR of the tsh locus. It was generated via a P-element swap that replaced the *p{GawB}* insertion of *tsh-GAL4* (Calleja et al., 1996) with *P{UpP65L}* and the enhancer-trap LexA construct. Proper targeting and orientation of *P{UpP65L}* were confirmed by splinkerette PCR and sequencing by J.-M. Knapp (unpublished results).

The empty stock *yw;;attP2* (Jenett et al., 2012) was also crossed to *UAS-CsChrimson; tsh-GAL80* as a baseline control. The following driver lines were used:

| Neuron | GAL4 driver | Figure | Reference |
| --- | --- | --- | --- |
| MBON-e1 | *GMR74B11* | 2 | (Saumweber et al., 2018) |

To test whether the response to the activation of the olfactory neuron Or42b requested functional Kenyon cells, we made the following constructs: 1) effector stock: *13XLexAop2-CsChrimson-mVenus* (in *attP18*, (Klapoetke et al., 2014); BDSC 55137) ; ; *UAS-Kir2.1::GFP* (gift from Y. Aso), crossed to 2) driver stock: *w+CS ; Or42b-LexA* (in *JK22C*, (Fishilevich & Vosshall, 2005) ; *GMR14H06-GAL4* (in *attP2*, (Jenett et al., 2012); BDSC 48667). The effector stock was crossed to the pan-neuronal driver *GMR57C10-GAL4* ((Jenett et al., 2012); BDSC 80962) to verify Kir2.1 was blocking neurons (no larvae hatched from the crosses); and to the empty line *y w;attP40;attP2* (Pfeiffer et al., 2010) to have a reference response to optogenetic activation of Or42b. A second reference response was obtained by crossing the driver stock to the line *13XLexAop2-CsChrimson-mVenus* (in *attP18; (Klapoetke et al., 2014)*; BDSC 55137).

To further characterize the properties of CN-4 and MBON-m1, we crossed the Split-GAL4 lines SS02108 (for CN-4) and SS02163 (for MBON-m1) to the following constructs:

- *UAS-Kir2.1* (Baines et al., 2001) to hyperpolarize CNs and test their contribution in navigation behavior in a gradient of ethyl acetate.
- *UAS-GtACR2* (gift from V. Jayaraman and A. Claridge-Chang; (Govorunova et al., 2015; Mohammad et al., 2017) to observe behavioral response to acute hyperpolarization of the CNs.
- *20XUAS-IVS-Syn21-opGCaMP6f p10* (Chen et al., 2013); *14H06-LexAp65* (in *JK22C*; (Jenett et al., 2012)); *LexAop-TNTe* (gift from J. Simpson, unpublished line; (Sweeney et al., 1995) to image CNs’ response to odors while silencing MB pathway. This stock was also crossed to pan-neuronal line *GMR57C10-LexA* to verify TNTe was silencing neurons (no larvae hatched from the crosses). In addition, we directly verified that KC output was indeed blocked by training the *SS02108* experimental cross in odor-sugar pairing and observing no immediate memory as compared to a control wild-type group (Extended Data Figure 7).
- *pJFRC22-10xUAS-IVS-myr::tdTomato* (in *su(Hw)attP8*; (Pfeiffer et al., 2010); BDSC 32223); *72F11-LexAp65* (in *JK22c*; (Jenett et al., 2012)); *20xUAS-GCamp6f 15.693* (in *attP2*, (Chen et al., 2013)), *13xLexAop2-CsChrimson-tdTomato* (in *vk00005*; (Klapoetke et al., 2014); BDSC 82183) to image CNs’ response to odors before and after aversive training. To verify that larvae resulting from such crosses were capable of learning, we trained first-instar larvae that resulted from a cross between *SS02108* and the above line, following a pairing protocol between odor and optogenetic activation. The larvae fed with retinal showed strong aversive short-term memory, whereas the larvae without retinal did not (see extended Fig. 5 for details).

### Learning experiments and odor navigation

Learning experiments were performed as previously described (Eichler et al., 2017; Eschbach et al., 2020; Saumweber et al., 2018). Briefly, two groups of 30 third-instar larvae were separated from food and underwent a training procedure involving odor and light exposures, either fully overlapping in time (paired group), or fully non-overlapping (unpaired group). The paired group was placed for 3 minutes on 4% agarose plates and exposed to constant red-light illumination (629 nm, 2.5 µW/mm2) paired with the presentation of 12 µl of odor ethyl-acetate 4.10^-6^ dilution in distilled water) absorbed on two filter papers located on the plate lid. These larvae were then transferred to a new plate with no odor and in the dark for 3 minutes. This paired training cycle was repeated three times in total. The unpaired group of larvae underwent odor presentation in the dark and red light without odor following the same protocol. These larvae were then immediately transferred to a 25 cm^2^ custom-made odor-delivery arena (described in (Gershow et al., 2012) covered with 4% agar and illuminated with infrared (for detection) and red (for memory expression, (Gerber & Hendel, 2006); 2.5 µW /mm2) light. A linear gradient of odor was generated by modulating the opening times of 32 3-way valves. One input to each valve was fed with odorized air (generated by bubbling air through a 4.10^-6^ dilution of EtAc in distilled water and the other with humidified air (generated by bubbling air through distilled water). The outlet of each valve projected into one of 32 parallel channels pushing humidified air across the chamber; a microcontroller and custom electronics were used to switch the valve opening times to produce a linear gradient (Gershow et al., 2012). The navigation behavior of the larvae was recorded with a camera (DALSA Falcon 4M30) and analyzed with machine vision developed in Matlab.

To test the role of CNs in innate odor navigation we placed larvae expressing Kir2.1 in neurons or control larvae in a gradient of odor. *Ca*. 30 larvae were separated from food, rinsed, and placed in an 25×25 cm square dish filled with 4% agar and whose one side of the lid was covered with 5 glued filter papers (7 mm^2^) each loaded with 12µl of 10^-4^ ethyl acetate. The navigation behavior of the larvae was recorded with a camera (DALSA Falcon 4M30) and analyzed with machine vision (Matlab).

For all navigation behaviors, bouts of individual trajectories were reconstructed offline, and interrupted when detection was compromised (e.g. if the larvae reached the border or crosses another larva’s path). To quantify the overall navigational response in the linear spatial gradient, we computed a navigation index by dividing the mean velocity of all larvae in the *x*-direction by the mean crawling speed. Hence, the navigational index was +/−1 if the larvae crawled uniformly straight up/down the gradient and 0 if the movement was unbiased. For each bout and at each time point, the position of the larva relative to the odor gradient was estimated as well as its likelihood to be engaged in a turn (details in (Gershow et al., 2012). To quantify turn-based navigation strategy, we compared the turn probability when the larvae were aligned towards the gradient (+/-15 degrees) to the turn probability when the larvae were aligned away from the gradient (+/-15 degrees). The experiments were repeated 7 to 10 times in each condition. The mean and s.e.m. of these parameters were computed for each experiment and further pooled for the repeats. A Welch z test was used for statistical comparisons.

### Optogenetic neural activation screen

Optogenetic activation experiments were performed as previously described (Ohyama et al., 2015). *Ca*. 30 larvae were separated from food by bathing them in a 20% sucrose solution for a maximum of 10 min. They were rinsed and placed into a square 25 cm^2^ behavior rig covered with 4% agar. We recorded videos of larval behavior, with a DALSA Falcon 4M30 camera for a total of 120 sec. At 30 and 75 sec, 15 sec-long pulses of red 660 nm red light (4 µW/mm2, Philips Lumileds) were applied. For statistics, speed (normalized to the baseline before the stimulations) and turn angle are averaged for all larvae and time windows corresponding to the two stimulations. For onset response, time window was taken during 5 sec. after light on for offset response, during 10 sec. after light off. Welch z tests were used for statistical comparisons.

### Circuit mapping and electron microscopy

We reconstructed neurons and annotated synapses in a single, complete central nervous system from a 6 hr old female *Canton S G1 x w1118* larva, acquired with serial section transmission EM at a resolution of 3.8 x 3.8 x 50 nm, that was first published along with the detailed sample preparation protocol (Ohyama et al., 2015). Briefly, the CNS was dissected and placed in 2% gluteraldehyde 0.1 M sodium cacodylate buffer (pH 7.4). An equal volume of 2% OsO_4_ was added and the larva was fixed with a Pelco BioWave microwave oven with 350-W, 375-W and 400-W pulses for 30 sec each, separated by 60-sec pauses, and followed by another round of microwaving but with 1% OsO_4_ solution in the same buffer. Next, samples were stained en bloc with 1% uranyl acetate in water and microwaved at 350 W for 3×3 30 sec with 60-sec pauses. Samples were dehydrated in an ethanol series, transferred to propylene oxide, and infiltrated and embedded with Epon resin. After sectioning the volume with a Leica UC6 ultramicrotome, sections were imaged semi-automatically with Leginon (Suloway et al., 2005) driving an FEI Spirit TEM (Hillsboro, OR), and then assembled with TrakEM2 (Cardona et al., 2012) using the elastic method (Saalfeld et al., 2012). The volume is available at https://l1em.catmaid.virtualflybrain.org/?pid=1.

To map the wiring diagram we used the web-based software CATMAID (Saalfeld et al., 2009), updated with a novel suite of neuron skeletonization and analysis tools (Schneider-Mizell et al., 2016), and applied the iterative reconstruction method (Schneider-Mizell et al., 2016). All annotated synapses in this wiring diagram fulfill the four following criteria of mature synapses (Ohyama et al., 2015; Schneider-Mizell et al., 2016) (1) There is a clearly visible T-bar or ribbon on at least two adjacent z-sections. (2) There are multiple vesicles immediately adjacent to the T-bar or ribbon. (3) There is a cleft between the presynaptic and the postsynaptic neurites, visible as a dark-light-dark parallel line. (4) There are postsynaptic densities, visible as dark staining at the cytoplasmic side of the postsynaptic membrane.

We validated the reconstructions as previously described (Ohyama et al., 2015; Schneider-Mizell et al., 2016), a method successfully employed in multiple studies (Berck et al., 2016; Fushiki et al., 2016; Goodman et al., 1981; Jovanic et al., 2016; Ohyama et al., 2015; Schneider-Mizell et al., 2016). Briefly, in Drosophila, as in other insects, the gross morphology of many neurons is stereotyped and individual neurons are uniquely identifiable based on morphology (Bate et al., 1981; Costa et al., 2016; Goodman et al., 1981). Furthermore, the nervous system in insects is largely bilaterally symmetric and homologous, with mirror-symmetric neurons reproducibly found on the left and the right side of the animal. We therefore validated neuron reconstructions by independently reconstructing synaptic partners of homologous neurons on the left and right side of the nervous system. With this approach, we have previously estimated the false positive rate of synaptic contact detection to be 0.0167 (1 error per 60 synaptic contacts) (Schneider-Mizell et al., 2016). Assuming the false positives are uncorrelated, for an n-synapse connection the probability that all n are wrong (and thus that the entire connection is a false positive) occurs at a rate of 0.0167^n^. Thus, the probability that a connection is a false positive reduces dramatically with the number of synaptic contacts contributing to that connection. Even for n = 2 synaptic contacts, the probability that a connection is not true is 0.00028 (once in every 3,586 two-synapse connections). We therefore consider ‘reliable’ connections those for which the connections between the left and right homologous neurons have at least 3 synapses each and their sum is at least 10. See (Ohyama et al., 2015; Schneider-Mizell et al., 2016) for more details. When predicting valence of CNs based on input from MBONs of known neurotransmitters and behavioral effects (approach or avoidance behaviors), we required a combined input of 5% from the appropriate MBONs to ensure that our predictions were robust.

### Similarity Matrices and Clustering

Adjacency matrices of synaptic connectivity were converted to binary connectivity matrices, representing only strong connections between hemilateral neuron pairs. A strong connection is defined as at least 3 synapses from the presynaptic left neuron and 3 synapses from the presynaptic right neuron onto the postsynaptic left and right neurons and a sum of at least 10 synapses total. Ipsilateral and contralateral connections are considered. Similarity is obtained by counting indices of value 1 that are observed at the same location in both the row neuron pair and the column neuron pair (matches) and counting the total number of value 1 indices that are only observed in the row or column alone, but not both (mismatches). The similarity score is the total number of matches, divided by the total number of matches and mismatches. Hierarchical clustering of similarity matrices was performed using R and heatmap.2 {gplots}.

### Identifying GAL4 lines that drive expression in MBONs and CNs

To identify GAL4 lines that drive expression in specific neurons, we performed single-cell FlpOut experiments (for FlpOut methodology see (Nern et al., 2015; Ohyama et al., 2015) of many candidate GAL4 lines (H. H. Li et al., 2014). We generated high-resolution confocal image stacks of individual neuron morphology (multiple examples per cell type). Most MBONs were uniquely identifiable based on the dendritic and axonal projection patterns (which MB compartment they project to and the shape of input or output arbor outside the MB). Some MBON pairs were too similar to be distinguished: MBON-h1/h2, g1/g2, and b1/b2.

### Brain explants imaging of CN-4 response to KC optogenetic activation

Central nervous systems were dissected in a cold buffer containing 103 mM NaCl, 3 mM KCl, 5 mM TES, 26 mM NaHCO_3_, 1 mM NaH_2_PO_4_, 8 mM trehalose, 10 mM glucose, 2 mM CaCl_2_, 4 mM MgCl_2_ and adhered to poly-L-lysine (SIGMA, P1524) coated cover glass in small Sylgard (Dow Corning) plates.

Optogenetic activation was done by red flood illumination (617nm High-Power Lightguide Coupled LED Source, Mightex) positioned above the sample. Light stimulations were performed with 1 or 15 sec duration and in 40 and 600 cycles of laser on/off pulses of 20 msec/5 msec. Each preparation underwent three types of light stimulation of increasing power: ca. 390 µW/mm^2^, 920 µW/mm^2^ and 4.6 mW/mm^2^. Only the data for the highest light power and longer duration is shown in Fig.4. The same stimulus was spaced with 30 sec for a total of three presentations in each scan. Each scan consisted in imaging CN-4 on a two-photon scanning microscope (Bruker) using a 60x 3 1.1 NA objective (Olympus). A mode-locked Ti:Sapphire laser (Chameleon Ultra II, Coherent) tuned to 925 nm was used for photo-activation of the GCaMP6f. Fluorescence was collected with photomultiplier tubes (Hamamatsu) after band-pass filtering. Images were acquired in line scanning mode (5.15 fps) for a single plane of the CNS.

For image analysis, image data were processed by Fiji software (Schindelin et al., 2012) and analyzed using custom code in Matlab (The Mathworks, Inc). Specifically, we manually determine the regions of interest (ROIs) from maximum intensity projection of entire time series images, and measure the mean intensity. In all cases, changes in fluorescence were calculated relative to baseline fluorescence levels (*F*_0_) as determined by averaging over a period of at least 2 sec. just before the optogenetic stimulation. The fluorescence values were calculated as (*F***_t_** -*F***_0_**)/*F***_0_**, where *F***_t_** is the fluorescent mean value of a ROI in a given frame. Analyses were performed on the average of the consecutive 3 stimulations and comparisons of before *vs*. after stimulation were done using a non-parametric Wilcoxon test for paired comparisons.

### Microfluidic device design

Odorant stimuli were delivered using a microfluidic device described in detail in (Si et al., 2019) and modified to deliver 3 odors instead of 13. The larva loading channel was 300 μm wide and 70 μm high, and tapered to a width of 60 μm in order to immobilize the larva. The tapered end was positioned perpendicular to a stimulus delivery channel to allowing odorant to flow past larval dorsal organ that houses 21 ORNs. The device was designed with a “shifting-flow strategy”, enabling odor changes without pressure or flow rate discontinuities (Chronis et al., 2007). An 8-channel device included two control channels located at the periphery, 3 odorant channels in the middle, and one water channel to remove odorant residue (the two remaining channels were blocked by a stopper). Each channel was of equal length to ensure equal resistance. During an experiment, a combination of three channels was always open: the water channel, one of the 3 odorant delivery channels, and one of the control channels. The 3 odorant channels could be sequentially opened to deliver any odorant. Switching between the two control channels directed either water or an odorant to flow past the larva’s ORNs.

Fluorescein dye was used to verify the spatial odorant profile in the device during stimulus delivery. The air pressure for stimulus delivery was set to 3 psi, where the switching time between water and odorant was estimated to be ∼20 ms.

The microfluidic device pattern was designed using AutoCAD. The design pattern was then transferred onto a silicon wafer using photolithography. The wafer was used to fabricate microfluidic devices using polydimethylsiloxane (PDMS) and following the standard soft lithography approach (Anderson et al., 2000). The resulting PDMS molds were cut and bonded to glass cover slips. Each microfluidic device was used for a few number of experiments and water- and air-cleaned between each of them in order to prevent contamination.

### Odorant delivery setup

Odorants were obtained from Sigma-Aldrich, diluted in deionized (DI) water (Millipore) and stored for no more than 3 days. We used n-amyl acetate (diluted in water for a 10^-3^ final concentration, AM), 3-octanol (10^-4^, OCT), ethyl acetate (10^-4^, EA), and methanol (10^-1^, ME). Each odorant concentration was stored in a separate glass bottle and delivered through its own syringe and tubing set. Panels of odorants were delivered using an 8-channel pinch valve perfusion system (AutoMate Scientific Inc.). Each syringe and tubing set contained a 30 mL luer lock glass syringe (VWR) connected to Tygon FEP-lined tubing (Cole-Parmer), which in turn was connected to silicone tubing (AutoMate Scientific Inc.). The silicone tubing was placed through the pinch valve region of the perfusion system and could allow for the passage or blockage of fluid flow to the microfluidics device. The silicone tubing was then connected to PTFE tubing (Cole-Parmer), which was then inserted into the microfluidic device. We used a DAQ board (National Instruments) to control the 8-channel pinch valve perfusion system using custom-written MATLAB code. This custom code allowed us to implement the on/off sequence of the valves and to synchronize valve control with the onset of recording in the imaging software (ScanImage).

During the entire recording, the larva experienced continuous fluid flow. The stimuli sequences consisted of five seconds of odorant pulses followed by a washout period using water.

### Calcium imaging

A first instar larva was loaded into a microfluidic device using a 1 mL syringe filled with 0.1% triton-water solution. Using the syringe, a larva was pushed towards the end of the channel, where the 60 μm wide opening mechanically trapped further larval movement. Each larva was positioned such that its dorsal organ (nose) was exposed to the stimulus delivery channel. Larvae were imaged at 35 fps using a multiphoton microscope equipped with a fast resonant galvo scan module (customized Bergamo Multiphoton, Thorlabs) controlled by ScanImage 2016 (http://www.scanimage.org). The light source was a femtosecond pulsed laser tuned to 925 nm (Mai Tai, Spectraphysics). The objective was a 25X water immersion objective (NA 1.1 and 2 mm WD, Nikon). The CNs neurites (dendrites and their contralateral axon terminals) were spanned in at least one brain hemisphere by a volume scan (6 slices with a step size of 2 μm).

For pairing of an odor with optogenetic activation of aversive neurons, a 660nm laser (Obis 660, Coherent) photostimulation (ca. 480 µW/mm^2^) was directed towards the terminals of the aversive neurons using a galvo-galvo module (Thorlabs) controlled by ScanImage software. The scans were usually not saved during the pairing period, as the imaging laser power were set to minimum power to avoid photobleaching associated with long-run recording. The CS+ was delivered for 20 sec, followed by a 20 sec laser stimulation which overlapped with CS+ for 17 sec. The CS- was presented alone for 20 sec. Each bout consisted in two CS+ and two CS- presentation; each interspaced by 50 sec of water flow, for a total of ca. 250 sec-long bout. The odors presented were different for different animals and were one of the following four combinations of CS: AM+/EA-, EA+/AM-, ME+/EA-, EA+/ME-. Eight consecutive pairing bouts were performed. The position of the larva in the channel was assessed between each bout and rectified with the triton-water-filled syringe if necessary. At the end of the eight bouts, the settings of the microscope were readjusted to allow optimal recording and the responses of the CN to delivery of the odors were reassessed the same way as before pairing. A single larva underwent between one and two of the pre-pairing and post-pairing scanning bouts.

The same system was used for co-stimulation of CNs with 10^-4^ ethyl acetate and optogenetic activation of Kenyon cells. Here two consecutive 5 sec-long odors and two consecutive 5 sec-long photo-stimulations were conveyed to the larva in a shuffled order, followed by two consecutive 5 sec-long joined delivery of odor and photo-stimulation. Each stimulation was interspaced with 20 +/-2 sec of water flow for a total of ca. 100 sec-long scanning bout. A single larva underwent between one and three of these bouts.

### Odor response analyses

The GCamp6f fluorescence (averaged intensity of z-projection) was calculated for a region of interest (CN’s neurite), subtracted to background intensity, and normalized to the tdtTomato signal emitted at CN’s membranes of the same region *F***_t_** = ((F*_GCamp__***_t_** / median(F*_GCamp_*)) *- F*_dTom_**t**_ / median(F*_GCamp_*). For each larva, one to two regions of interest (corresponding to the left and right hemispheres) were selected (by thresholding the projected maximum intensity image) for each larva and their fluorescence was averaged. 2 to 4 repetitive stimulations were averaged as well. Movement artifacts were corrected by aligning frames using the strongest signal (tdtTomato- or GCamp6f-derived) labelling the CN neurites and a combination of cross-correlation on Matlab (normxcorr2_general, ©2010, Dirk Padfield) and manual correction.

Changes in fluorescence were quantified as (*F***_t_** -*F***_0_**)/*F***_0_**, where *F*_0_ was the average fluorescence intensity sampled from the frames of the 5 sec preceding a stimulation. Quantifications of normalized mean and peak were the normalized value of, respectively, mean and maximum intensity for the frames during the stimulation (ON response) or for the 8 sec following the stimulation (OFF response). Statistical comparisons were done using a non-parametric Wilcoxon test for paired comparisons.

### Data exclusion

When movements artefacts were too important and rendered alignment impossible for a substantial part of the recording (ca. 10%), the data for this larva was discarded. Data for trained larvae was excluded if no calcium response was observed to any odors before and/or after the training session, as it likely indicated that the larva was dead or that the external sensory organs were not exposed to the odor flow.

## Acknowledgements

We thank Fly Light at HHMI Janelia Research Campus (JRC) for generating confocal images of the GAL4 lines, V. Jayaraman for sharing unpublished versions of CsChrimson and GCamp6f, K. Hibbard and JRC FLY Core for generating some of the fly stocks, Fly EM at JRC for generating the EM volume. We thank Feng Li (9%), Avinash Khandelwal (8%), Jennifer Lovick (5%), Ivan Larderet (5%), Timo Saumweber (5%), Volker Hartenstein (4%), Nadia Riebli (3%), Larisa Maier (2%), Alex Bates (2%), Laura Herren (1%), and many others (3%) for reconstructing arbor cable and synapses. Z. Zavala-Ruiz and the JRC Visiting Scientist Program and HHMI JRC for funding. C. E. and M. W. were supported by the ERC consolidator grant RG95162. M. Z. was also supported by the ERC consolidator grant RG95162 and Wellcome Trust Investigator Award RG86459. A. C.

## Author contributions

C. E. conceived the study, designed and performed experiments and data analysis, and wrote the manuscript. A. F. and M. W. reconstructed neuronal arbors and synapses from volume electron microscopy and analyzed the resulting anatomically-reconstructed neural circuits. A. F. identified neuron-specific GAL4 lines. M. W. performed data analysis and generated connectomics figures. M. Z. and A. C. conceived the study, performed data analysis, and wrote the manuscript. I. A. and J. V.-A. performed circuit reconstruction and analysis. B. A. and G. S. contributed experimental design and preliminary imaging data, B. T. C., R. G. and K. E. performed behavioral experiments. A. S. and M. G. contributed data analysis and experimental design. R. D. F., G. S.X.E. J. and J. W. T. contributed data and reagents.

## Extended Data Figure

**Extended Data Figure 1.**
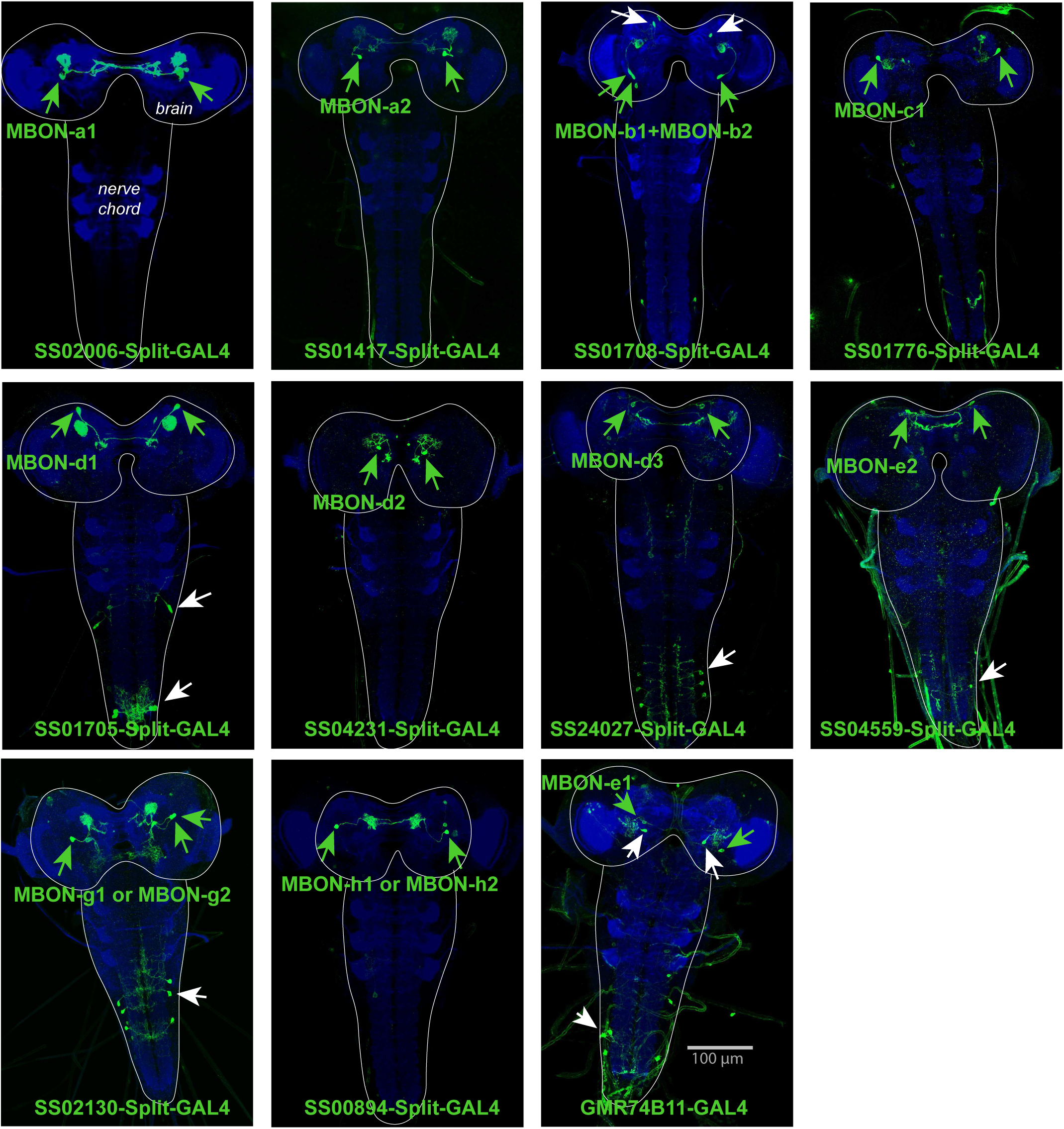
(related to Figure 2): Expression patterns of Split-GAL4 lines. Each panel shows a confocal maximum intensity projection of the complete CNS of third-instar larvae (outlined by the dotted line), with the neuropil labeled with anti-N-Cad antibody (blue) and the Split-GAL4 line expression pattern revealed by driving *UAS-myr-GFP* (green). Arrowheads indicate cell bodies of identified neurons: in green the targeted neuron, in white other neurons. The Split-GAL4 lines *SS02006* (for MBON-a1), *SS01417* (for MBON-a2), *SS01776* (for MBON-c1), *SS04321* (for MBON-d2), *SS00894* (for MBON-h1 and -h2) drive expression specifically in the neurons of interest. The lines *SS01708* (for MBONb1 and -b2) also drive expression in another brain neuron type. *SS01705* (for MBON-d1), *SS24027* (for MBON-d3), *SS04559* (for MBON-e2), *SS02130* (for MBON-g1 or -g2) target interneurons in the ventral nerve cord in addition to the neuron of interest; and *GMR-74B11* (MBON-e1) drive expression in another brain neuron and in neurons of the ventral nerve cord (eliminated using the GAL80 repressor in *teashirt-LexA* for the ventral nerve cord).

**Extended Data Figure 2.**
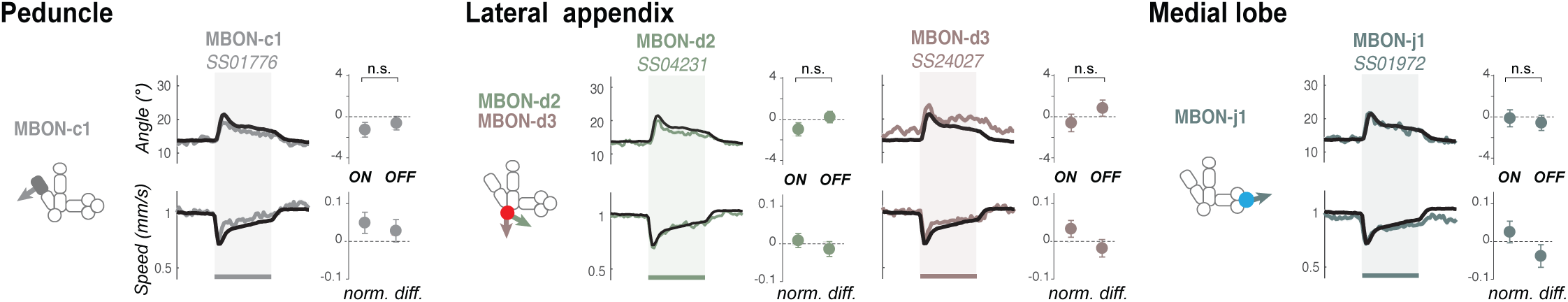
(related to Figure 2): Some MBONs do not promote visible response when optogenetically activated. The behavior of larvae expressing CsChrimson in one or two pairs of MBONs were recorded and compared to the response of the control empty driver line (shown in *black*, itself shows a slight increase in turn in response to light onset). All experiments include at least 6 runs with *ca*. 40 animals per run. Here we show the responses evoked by the lines driving expression in MBONs and that were not distinguishable from the control response. The name of the Split-GAL4 line used is shown in italic, below the neuron targeted by the line. The lines that drove expression in MBONs with visible activation phenotypes (approach- or avoidance-like) are shown in Fig. 2. The schematics depict the compartments where the MBONs extend their dendrites, filled with color indicating which type of memory this compartment can host (Eschbach *et al*., 2020): appetitive short-term (*blue*), aversive short-term (*red*), or unknown (*grey*). Plots are as in Fig. 2 and show mean +/- s.e.m for turn angle and speed over time (*left*) and normalized over baseline and control line for the 5 sec-long time window following light onset or offset (*right*). n.s.: p>0.05 of difference to control (Welch Z test).

**Extended Data Figure 3.**
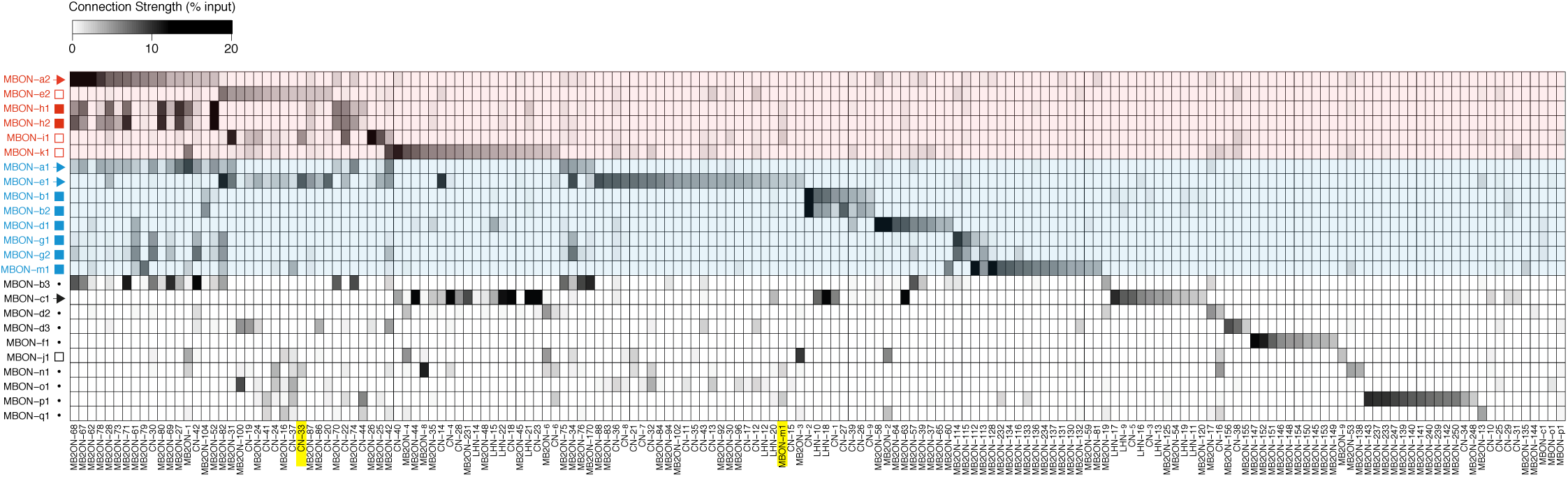
(related to Figure 3): Connectivity Matrix. Connectivity matrix showing normalized synaptic input (expressed as % input) each MB2ON (*columns*) receives from each MBON (*rows*). MBONs are color-coded according to the type of behavior they may promote (Fig. 2b-g,i): approach (*blue*), avoidance (*red*), or undefined (*black*). The symbols indicate the type of neurotransmitter they release when known from previous immunostaining experiments (Eichler *et al*., 2017). Colored shades indicate the type of valence code carried by the MBON onto the MB2ON: *light blue,* positive valence (carried either by excitatory approach-promoting MBON or by inhibitory avoidance-promoting MBON), *light red*, negative valence (carried either by excitatory avoidance-promoting MBON or by inhibitory approach-promoting MBON). Yellow boxes highlight neurons in which experiments were performed, namely CN-33 and MBON-m1. Normalized synaptic input (% input) is calculated by number of synapses per cell, divided by total number of postsynaptic sites in the column neuron.

**Extended Data Figure 4.**
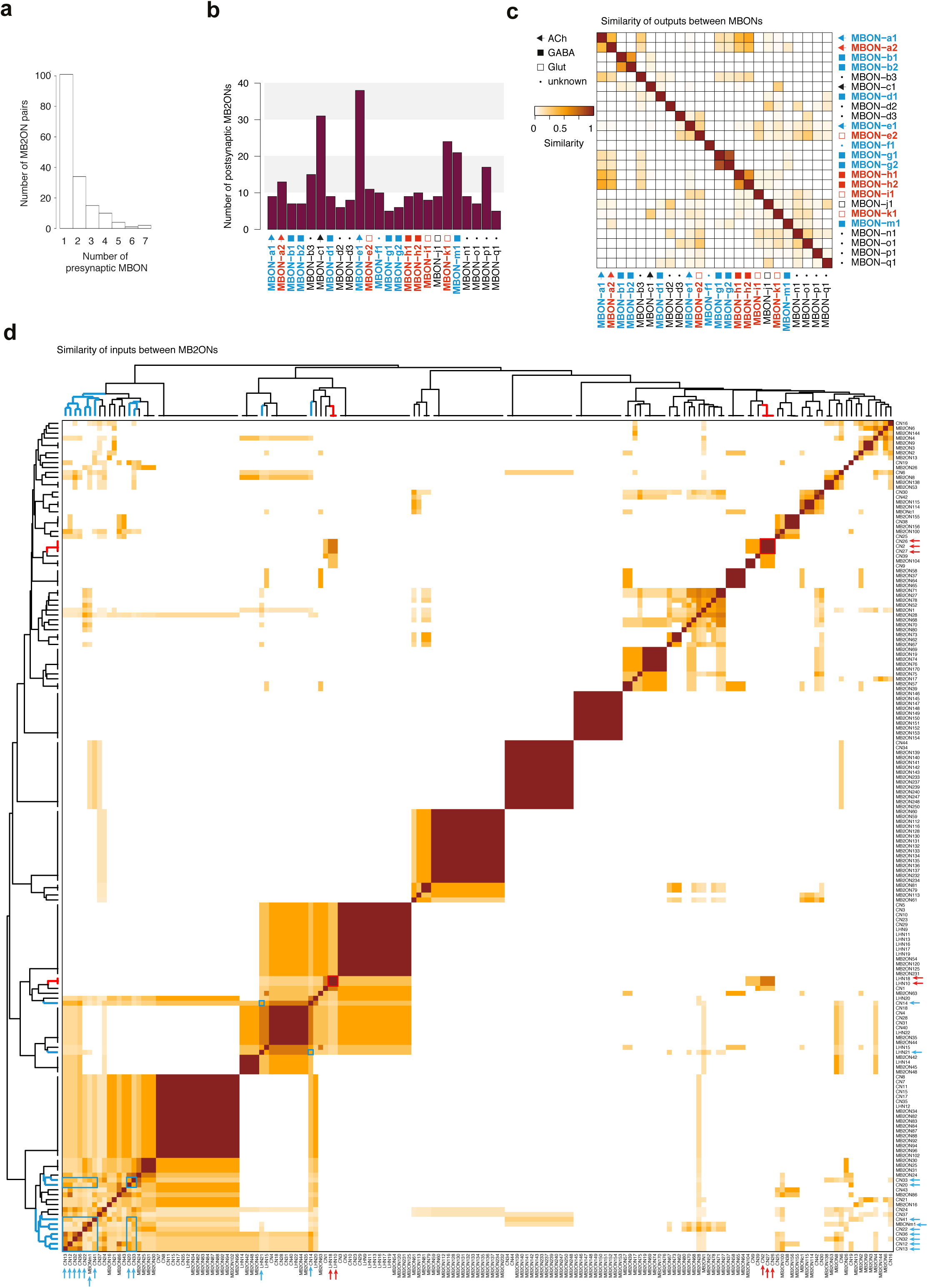
(related to Figure 3): Analysis of M2BONs inputs. For all the analyses (**a-d**), left-right homologous neurons are pooled. **a.** Histogram of number of strong presynaptic connections from MBONs to each MB2ON. **b.** Number of strong postsynaptic connections in MB2ONs from each MBON. **c.** Matrices of similarity between MBONs based on their total number of outputs onto MB2ONs. Similarity is obtained by counting the total number of outputs in a row neuron that are also outputs in a column neuron (matches) and counting the total number of outputs that are only observed in row or column neurons, but not both (mismatches). The similarity score is the total number of matches, divided by the total number of matches and mismatches. An output connection is only considered if there are at least 3 synapses from the presynaptic left neuron and 3 synapses from the presynaptic right neuron onto the postsynaptic left and right neurons and a sum of at least 10 synapses total. Ipsilateral and contralateral connections are considered. Most MBONs have a unique combination of postsynaptic partners and display low similarity scores. **d.** Similarity matrix of MB2ONs based on their total number of inputs from MBONs. Methodology similar to c., but comparing column neuron inputs, instead of row neuron outputs. Many MB2ONs have low similarity scores but there are some groups that receive strong connections from the same MBON(s). Hierarchical clustering was applied to the similarity scores to sort MB2ONs. Blue and red arrows indicate previously extrapolated positive and negative valence neurons, respectively. Blue and red boxes highlight similarity between positive and negative valence neurons, respectively, within clusters.

**Extended Data Figure 5.**
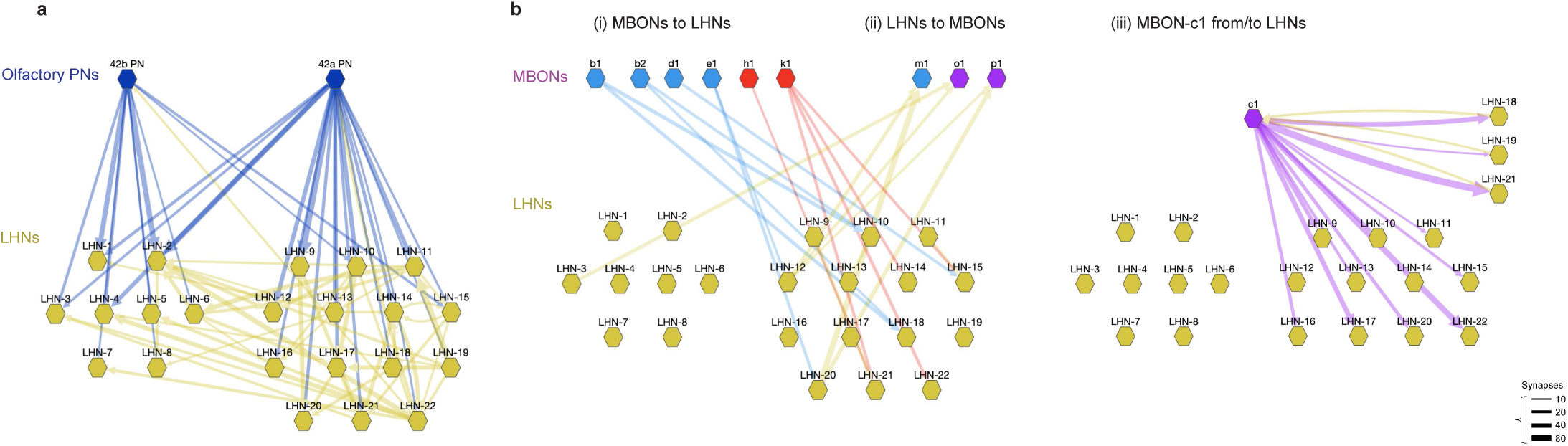
(related to Figure 3): Circuit diagram of MBONs and LHNs. **a.** The EM circuit graph of the 22 LHNs downstream of PN42a and PN42b olfactory projection neurons (likely tuned to attractive odors). The network reveals many common targets for the two types of PNs and tight interconnection between these putatively appetitive LHNs. Each node represents a pair of left-right homologous neurons. Connections with less than 10 synapses are not included. The left cluster of LHNs 1-8 is not postsynaptic of MBONs, whereas the right cluster of LHNs 9-22 is. **b.** Detailed EM circuit graph of MBONs and LH neurons: (i) synaptic connections from MBONs to LHNs (*blue* and *red* arrows; positive and negative MBON valence, respectively), (ii) synaptic connections from LHNs to MBONs (*yellow* arrows), (iii) MBON-c1 from/to LHNs. Unlike other MBONs, MBON-c1 has dense connections with many LHNs (*yellow* arrows from LHNs, *purple* arrows to LHNs); it also receives connections from multiple neurons, and only axo-axonic inputs from a few MBONs (MBON-d1, -g1 and - g2). Because of this specific type of connectivity, its function will not be discussed further in this article. Hierarchical clustering was applied to the similarity scores to sort CNs.

**Extended Data Figure 6.**
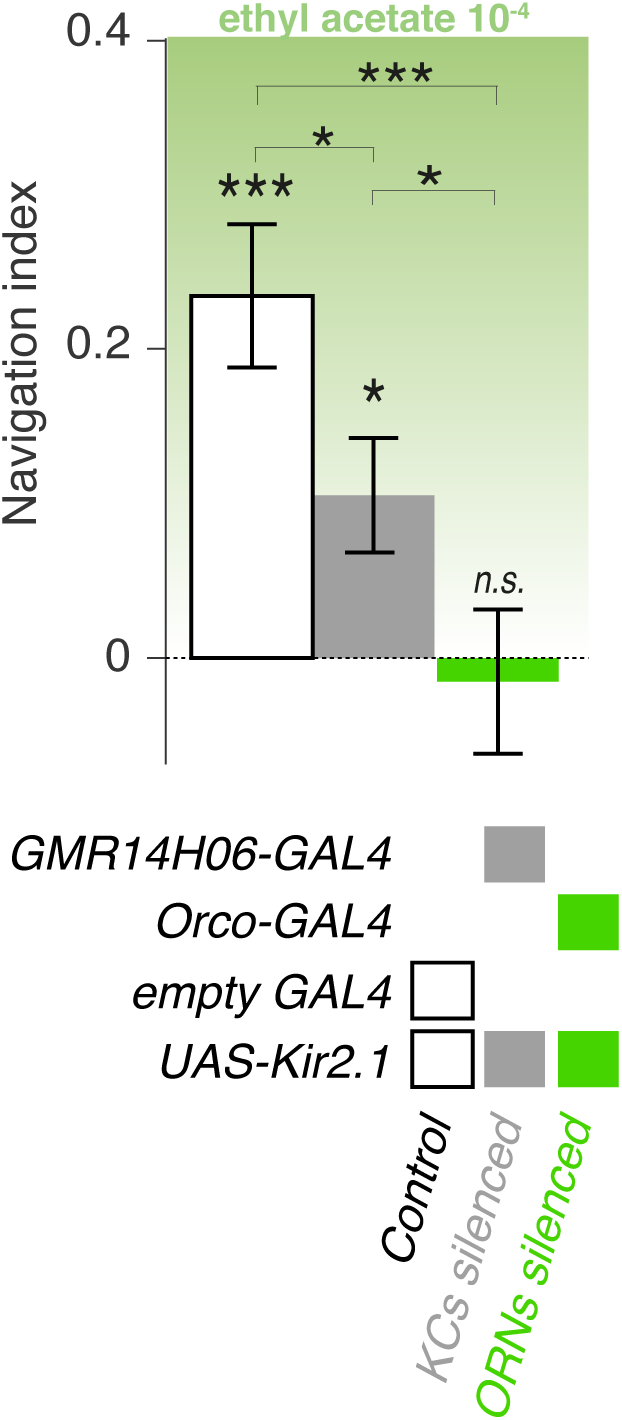
(related to Figure 3): Functioning Kenyon Cells are required for normal approach behavior of an odor. **a.** The behavior of larvae in a linear gradient of odor (ethyl acetate) is assessed in naive larvae with silenced Kenyon Cells (using GMR14H06-GAL4 crossed to UAS-Kir2.1). The navigational indices in larvae with non functioning MBs are lower than the navigational indices in the control larvae (empty GAL4 crossed to UAS-Kir2.1). This indicates that the MB can participate in innate attraction to an odor, consistent with results described in Fig. 3e.

**Extended Data Figure 7.**
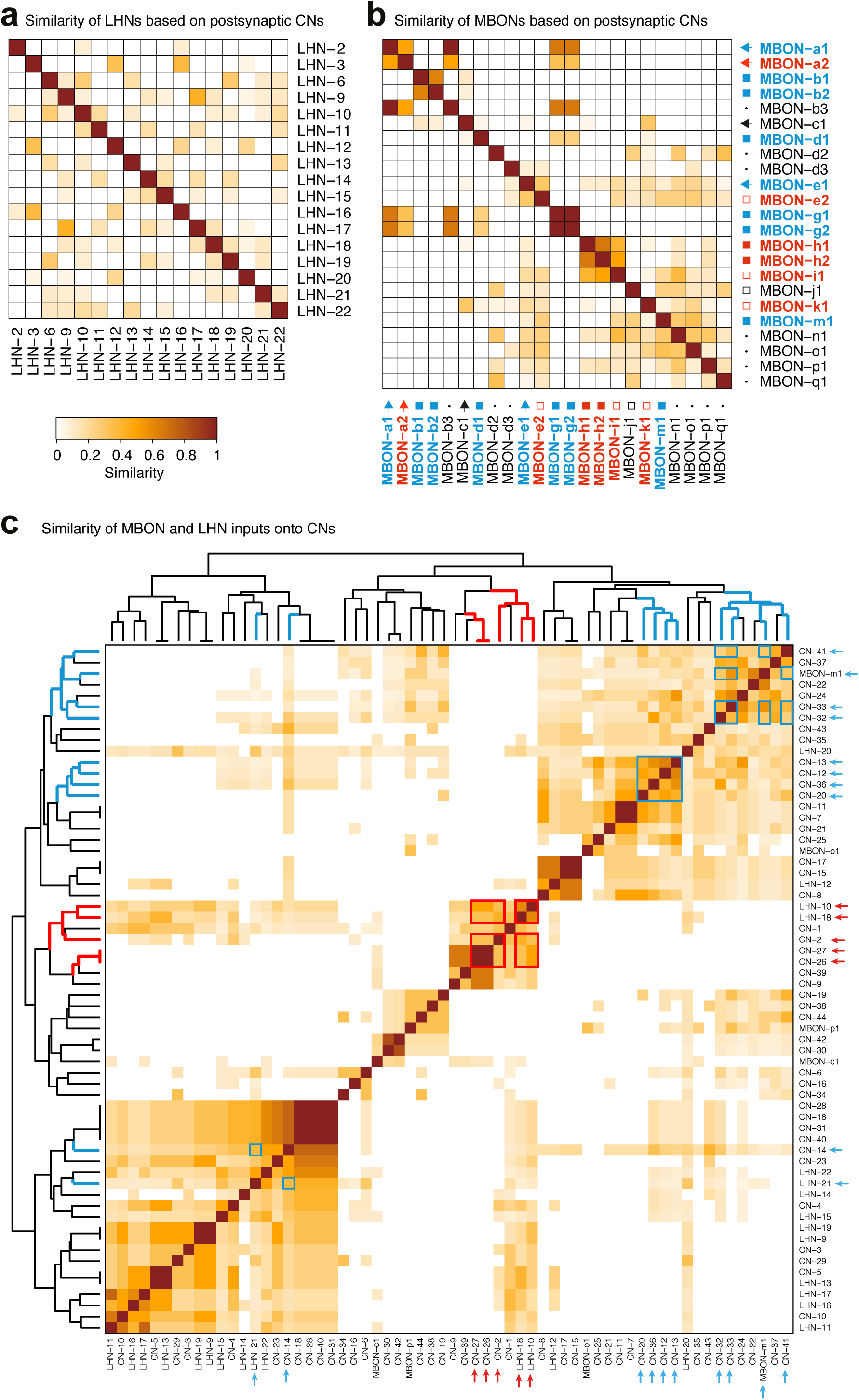
(related to Figure 3): Analysis of CNs inputs. **a-b.** Matrices of similarity between neurons based on their total number of strong output connections onto CNs (**a-b**), defined as the subset of MB2ONs which receives inputs from both the LH pathway downstream of ORN42a/ORN42b and MBONs. Similarity is obtained following the method described in Extended Data Figure 3. **a.** Similarity matrix of LHNs based on outputs to CNs. **b.** Similarity matrix of MBONs based on outputs to CNs. Most MBONs and LH neurons have a unique combination of postsynaptic CN partners. **c.** Similarity matrix of CNs based on their total number of inputs from MBONs and LHNs. Most CNs receive a unique combination of MBONs and LHNs inputs. Hierarchical clustering was applied to the similarity scores to sort MB2ONs. Blue and red arrows indicate previously extrapolated positive and negative valence neurons, respectively. Blue and red boxes highlight similarity between positive and negative valence neurons, respectively, within clusters.

**Extended Data Figure 8.**
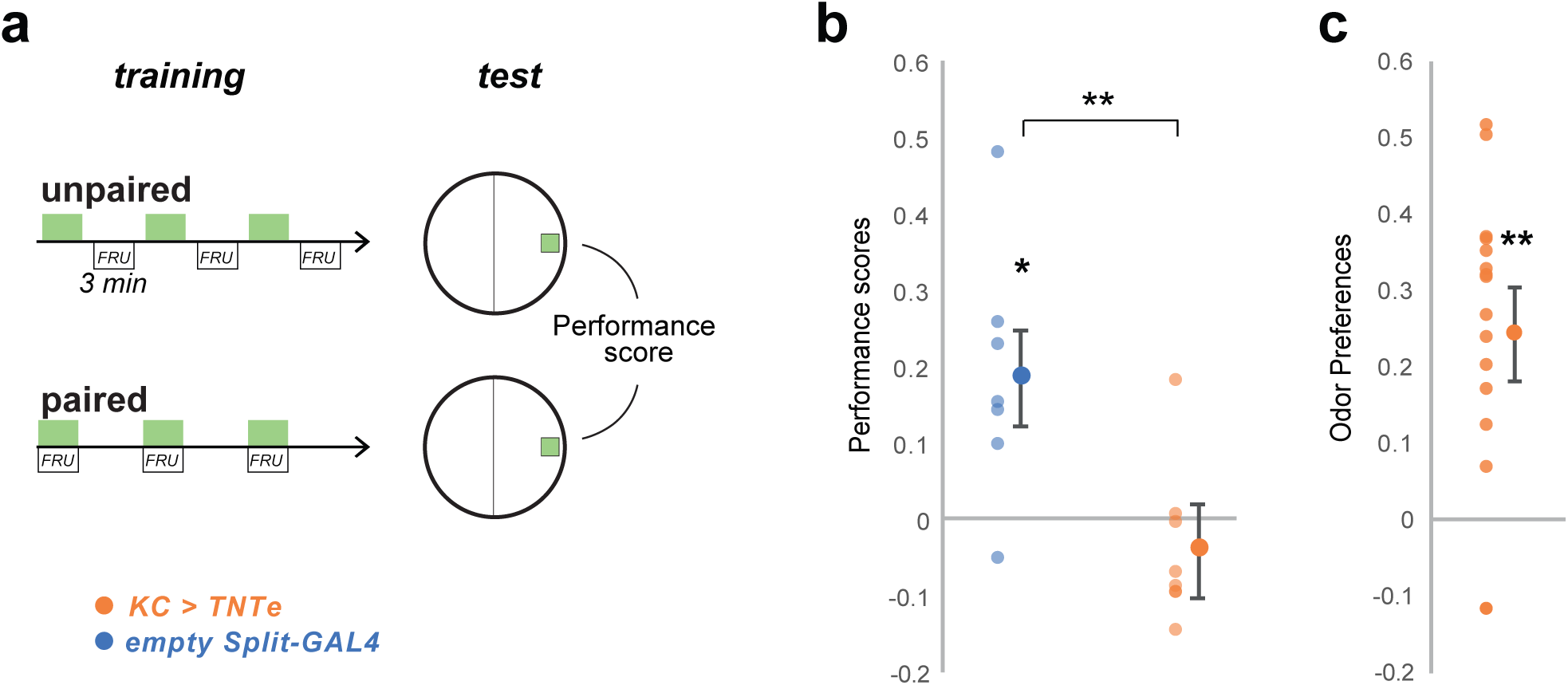
(related to Figure 4): Silencing KC impairs olfactory memory performances but maintains olfactory perception. We verified that the larvae expressing TNTe in KC were impaired in associative learning. **a.** We trained groups of 30 third-instar larvae in sets of two. For each pair, one group, the “paired group”, was presented with ethyl acetate (green rectangles) and fructose-supplemented agar for 3 times 3 min-long pairing intercalated with 3 min of no odor and pure agar. The other group, the “unpaired group”, received ethyl acetate for 3 min and fructose-supplemented agar for the 3 next min, 3 times with no overlapping. The two groups were then tested for their preference for ethyl acetate, which was estimated by Pref_EA_ = (N_EA_ – N_air_) / (N_EA_ + N_air_), and a Performance Score was computed by subtracting the Pref_EA_ in the “paired” group to the Pref_EA_ obtained in the “unpaired” group. A positive score indicates appetitive memory, whereas a zero score indicates no memory. **b.** The third-instar larvae of the same genotype as in Fig.6b (KC>TNTe ; CN-33>GCamp6f, N=8) did not show appetitive short-term memory while the control line (empty Split-GAL4, N=7) did. *: p < 0.05, **: p < 0.01, Wilcoxon test. Individual datapoints and mean +/- s.e.m. are shown. **c.** After either the paired or unpaired training procedure, the experimental larvae (KC>TNTe ; CN-33>GCamp6f, N=8) all exhibited attraction to the trained odor, indicating that memory but not olfactory behavior was not affected. Statistics are the same as in **b**.

**Extended Data Figure 9.**
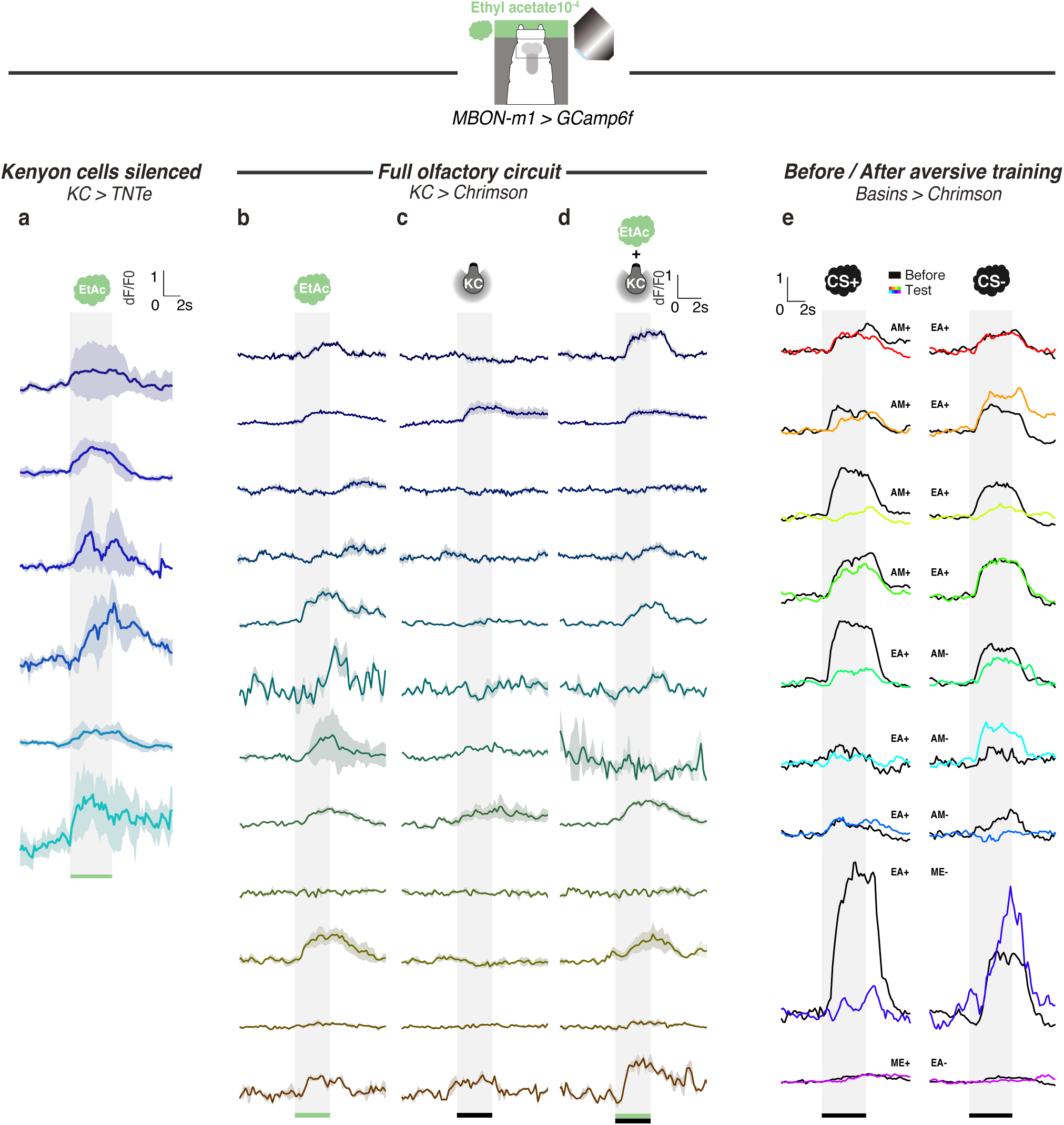
(related to Figures 4): Calcium responses of MBON-m1 for each individual. Calcium activity of MBON-m1 was imaged *in vivo* (using *SS02163-GAL4 > UAS-GCamp6f*) in larvae trapped in microfluidic device (Si *et al.*, 2019) and exposed to the odor ethyl acetate (**a**,**b**,**d**) and/or to optogenetic activation of Chrimson-expressing KCs (**c**,**d**). Each curve shows fluorescence normalized to baseline (before odor presentation) and averaged over 2 to 4 repeats. Data in **b-d** are from the same animals, as indicated by the same color. **a.** Response of MBON-m1 to ethyl acetate in larvae with silenced MB (using *14H06-LexA > LexAop-TNTe*). **b.** Response of MBON-m1 to ethyl acetate in larvae with intact MB. **c.** Response of MBON-m1 to activation of MB (using *14H06-LexA > LexAop-CsChrimson*). **d.** Response of MBON-m1 to coincident odor delivery and activation of MB. **e.** Response of MBON-m1 to two odors, CS+ and CS-, before and after the CS+ was paired with optogenetic activation of the nociceptive neurons Basins (olfactory aversive training).

**Extended Data Figure 10.**
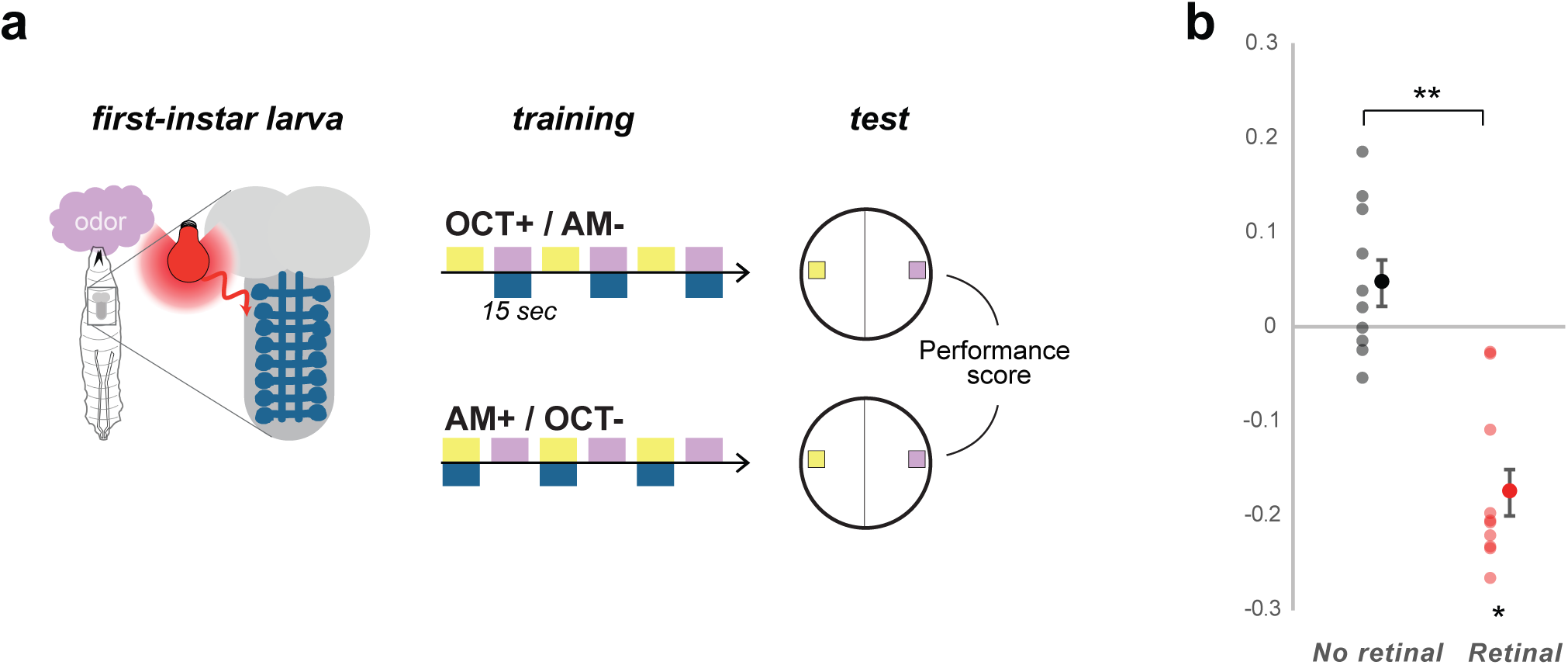
(related to Figure 5): Learning performances in first-instar larvae. We verified that the larvae we imaged in Fig.5 were capable of associative learning. **a.** To do this, we trained pairs of groups of 30 first-instar larvae to associate either 3-octanol (OCT+) or n-amyl acetate (AM+) with optogenetic activation of the aversive Basins neurons (drawn in dark blue) for 15 sec long pairing whereas the other odor was presented alone for 15 sec. The larvae were then tested for their choice between the two odors, which was estimated by Pref_AM_ = (N_AM_ – N_OCT_) / (N_AM_ + N_OCT_), and a Performance Score was computed by subtracting the Pref_AM_ after OCT+/AM- training to the Pref_AM_ obtained in the group that received AM+/OCT- training. A negative score indicates aversive memory, whereas a zero score indicates no memory. **b.** The first-instar larvae of the same genotype as in Fig.5 form an aversive short-term memory if they were raised in the presence of trans-retinal in the food (necessary for a functional channelrhodopsin and thus for the aversive neurons to be depolarized during training). N=10, **: p = 0.002, Wilcoxon test.

**Extended Data Figure 11.**
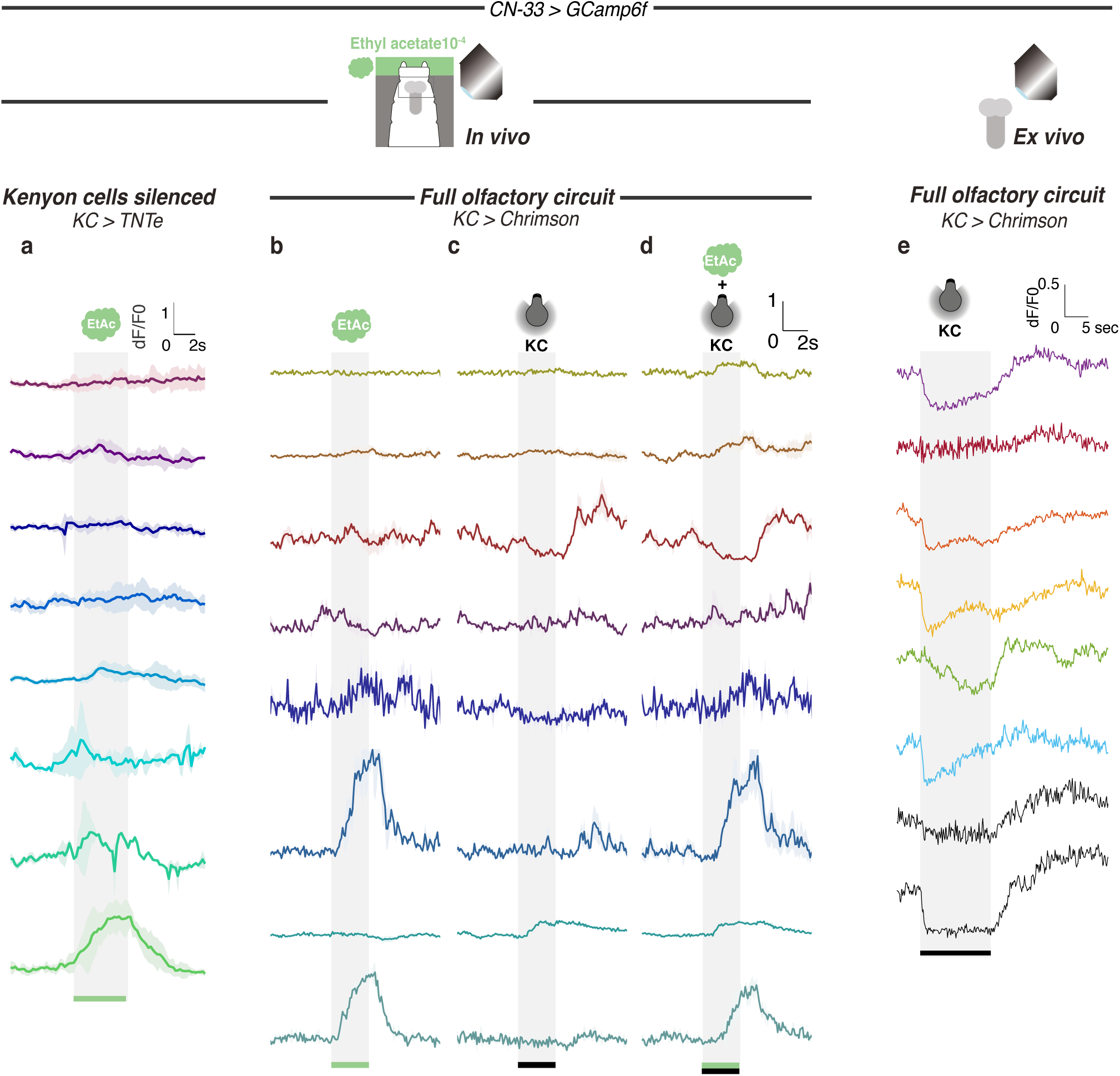
(related to Figures 6): Calcium responses of CN-33 for each individual. Calcium activity of CN-33 was imaged *in vivo* (using *SS02108-GAL4 > UAS-GCamp6f*) in larvae trapped in microfluidic device (Si *et al.*, 2019) and exposed to the odor ethyl acetate (**a**,**b**,**d**) and/or to optogenetic activation of Chrimson-expressing KCs (**c**,**d**). Each curve shows fluorescence normalized to baseline (before odor presentation) and averaged over 2 to 4 repeats. Data in **b-d** are from the same animals, as indicated by the same color. **a.** Response of CN-33 to ethyl acetate in larvae with silenced MB (using *14H06-LexA > LexAop-TNTe*). **b.** Response of CN-33 to ethyl acetate in larvae with intact MB. **c.** Response of CN-33 to activation of MB (using *14H06-LexA > LexAop-CsChrimson*). **d.** Response of CN-33 to coincident odor delivery and activation of MB. **e.** Response of CN-33 to activation of MB in brain explants.

**Extended Data Figure 12.**
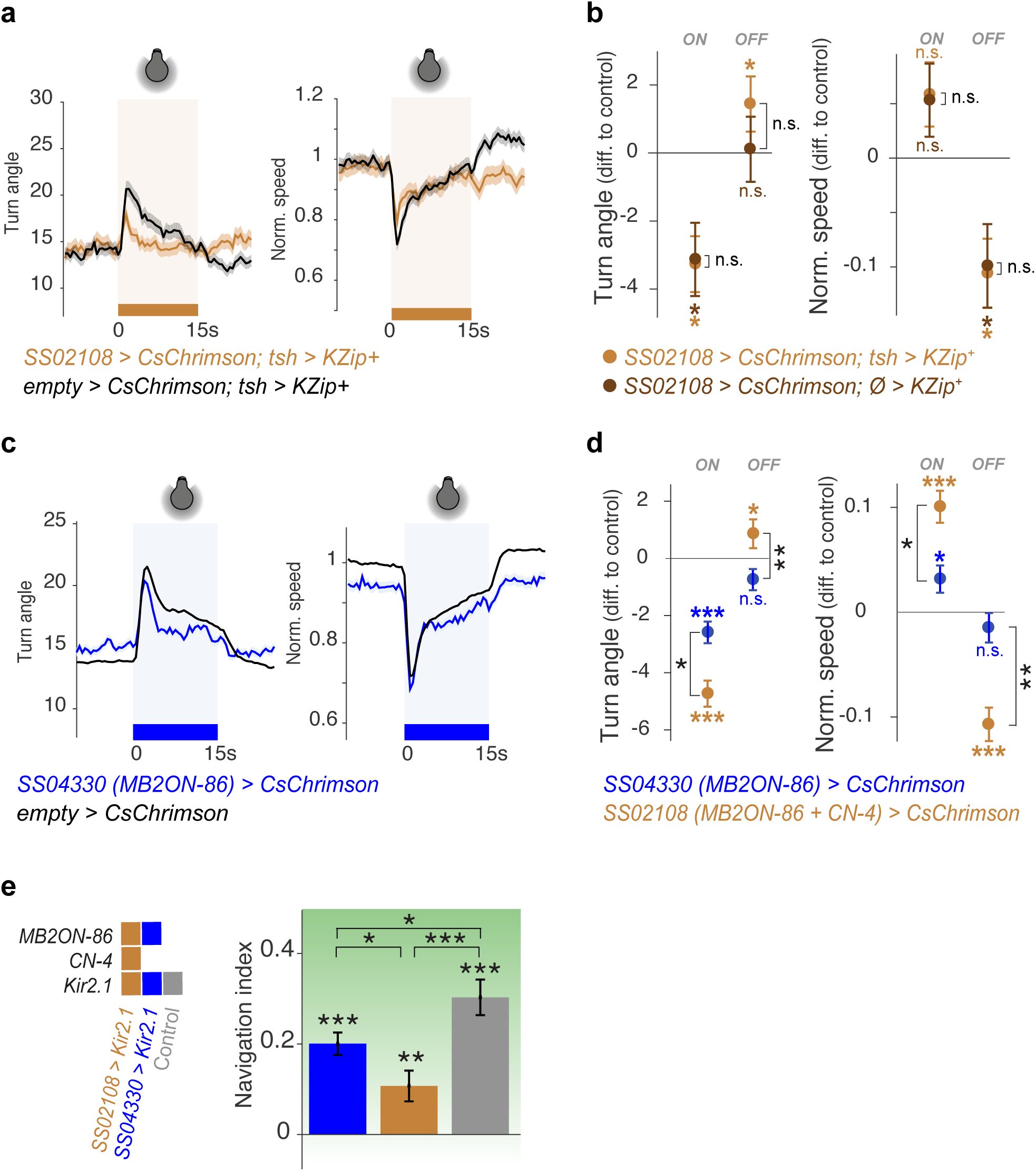
(related to Figure 6): Control crosses in activation or silencing experiments confirm that CN-33 is involved in approach behavior. **a.** We removed *SS02108*-driven neural expression in the nerve cord by expressing the Split-GAL4 repressor Killer Zipper (*LexA_op_-KZip^+^::3xHA*, Dolan *et al*., 2017) under the control of *teashirt-LexA* driver (*SS02108>CsChrimson; tsh>KZip^+^*). With this genetic manipulation we verified that the approach behavior of larvae observed in Fig. 4b was due to the brain neurons present in the *SS02108* expression pattern. **b.** Removing neural expression in the nerve cord (with *SS02108>CsChrimson; tsh>KZip^+^*) did not change optogenetically induced behavior as compared to not removing it (with *SS02108>CsChrimson; Ø>KZip^+^*). N_animals_= 378,246, *: p<0.05, Welch Z test. **c.** The line *SS04330* drives expression in MB2ON-86 alone (Eschbach *et al.,* 2020) and is thus used to verify how specific to CN-33 are the behavioral effects found with the line *SS02108* (which targets both CN-33 and more weakly MB2ON-86). Of note *SS04330* did not drive expression at first instar stage and could thus not be used as a control for imaging. Optogenetic activation of MB2ON-86 alone evoked turn decrease at the onset of the light, consistent with an approach-like behavior. However, it did not induce an increase of turns at the light offset. **d.** As shown in Fig. 4b, the joined activation of CN-33 and MB2ON-86 did induce both a strong onset approach-like response, and an offset approach-like response. Here, only the offset behavioral component was found to be significantly different between the two lines. Thus, at least the offset part of the response observed with *SS02108* is due to CN-33. N= 892 (*SS04330*), 657 (*SS02108*), *: p<0.05, **: p<0.001, Welch Z test. **e.** Constitutive silencing of MB2ON-86 affects navigation performance. N_animals_= 135,92,211, N_experiments_= 7,4,6; *: p<0.05, **: p<0.001, ***: p<0.001, Welch Z test.

